# The Hox transcription factor Ultrabithorax binds RNA and regulates co-transcriptional splicing through an interplay with RNA polymerase II

**DOI:** 10.1101/2021.03.25.434787

**Authors:** Julie Carnesecchi, Panagiotis Boumpas, Patrick van Nierop y Sanchez, Katrin Domsch, Hugo Daniel Pinto, Pedro Borges Pinto, Ingrid Lohmann

**Affiliations:** Heidelberg University, Centre for Organismal Studies (COS) Heidelberg, Department of Developmental Biology, Heidelberg, Germany; Friedrich-Alexander-University Erlangen-Nürnberg, Department Biology, Division of Developmental Biology, Erlangen, Germany; Department of Cell Biology, Albert Einstein College of Medicine, New York, NY, USA

**Keywords:** Hox, mRNA splicing, transcription, RNA, RNA Polymerase II, homeodomain, Drosophila

## Abstract

Transcription Factors (TFs) play a pivotal role in cell fate decision by coordinating distinct gene expression programs. Although most TFs act at the DNA regulatory layer, few TFs can bind RNA and modulate mRNA splicing. Yet, the mechanistic cues underlying TFs function in splicing remain elusive. Focusing on the *Drosophila* Hox TF Ultrabithorax (Ubx), our work shed light on a novel layer of Ubx function at the RNA level. Transcriptome and genome-wide binding profiles in embryonic mesoderm and *Drosophila* cells indicate that Ubx regulates mRNA expression and splicing to promote distinct functions in defined cellular contexts. Ubx modulates splicing via its DNA-binding domain, the Homeodomain (HD). Our results demonstrate a new RNA-binding ability of Ubx in cells and *in vitro*. Notably, the N51 amino acid of the HD, which mediates Ubx-DNA interaction, is non-essential for Ubx-RNA interaction *in vitro* but is required *in vivo*. We find that the N51 amino acid is necessary to mediate interaction between Ubx and the active form of the RNA Polymerase II (Pol II S2Phos) in *Drosophila* cells. By combining molecular and imaging approaches, our results reveal that Ubx mediates elongation-coupled splicing via a dynamic interplay with active Pol II and chromatin binding. Overall, our work uncovered a novel role of the Hox TFs at the mRNA regulatory layer. This could be an essential function for other classes of TFs to control cell diversity.

## INTRODUCTION

Eukaryotic gene expression is a fascinating process: it creates various proteomes from limited genetic material, thereby promoting cell and tissue diversity in multicellular organisms. To do so, genes are transcribed into pre-mRNAs that undergo a series of RNA processing events such as 5’capping, splicing and 3’polyadenylation (Moore and Proudfoot, 2009). Splicing, the mechanism of exon/intron retention or excision, plays an important role in proteome diversity (Chen and Manley, 2009). It relies on the dynamic assembly of a ribonucleoprotein complex termed spliceosome and numerous accessory proteins that impel the RNA fate by selecting the appropriate splice sites (Wahl et al., 2009). Splicing is regulated at several levels in the nucleus (Bentley, 2014; Chen and Manley, 2009; Galganski et al., 2017; Spector and Lamond, 2011; Wahl et al., 2009). Notably, it mainly happens co-transcriptionally (Blencowe, 2006; Hegele et al., 2012; Wahl et al., 2009) and depends on the RNA Polymerase II (Pol II) elongation activity (Bentley, 2014; Carrocci and Neugebauer, 2019; Khodor et al., 2011; de la Mata et al., 2003; Oesterreich et al., 2011; Saldi et al., 2016). Remarkably, a single pre-mRNA can be spliced in different ways thereby diversifying transcript isoforms and proteins (Blencowe, 2006). This unique strategy termed Alternative Splicing (AS) contributes greatly to the diversity of cell and tissue identities in complex organisms (Auboeuf, 2018). Despite its fundamental contribution to the proteome diversity (Auboeuf, 2018; Moore and Proudfoot, 2009), realising how the transcriptional and splicing programs are coordinated is still a challenge in Biology.

Transcription factors (TFs) are key players of gene expression. They coordinate specific yet flexible transcriptional programs thereby orchestrating developmental diversity and tissue maintenance of living organisms (Junion et al., 2012; Rhee et al., 2014). They do so by recognizing DNA-binding sites and regulating target genes in a precise spatial and temporal manner. Intriguingly, most TFs act at the DNA regulatory layer, however few can bind RNA and/or modulate mRNA splicing (Auboeuf, 2002; Girardot et al., 2018; Han et al., 2017; Rambout et al., 2018). Notably, the TF Sox9 regulates gene expression via distinct functions on transcription and splicing (Girardot et al., 2018). The homeodomain (HD) containing TF Bicoid (Bcd) has been shown to interact with *caudal* RNA in *Drosophila* thereby inhibiting translation (Rödel et al., 2013). Other TF families such as the C2H2-Zinc-finger and the hormone nuclear receptors can modulate splicing, the latter via cofactors interaction (Auboeuf, 2002; Cramer et al., 1997). Despite the strong evidences supporting that TFs could be major regulators of mRNA splicing, the mechanistic cues underlying TFs function in AS remain elusive. More importantly, how the mRNA-regulatory function of TFs impacts on cell fate decisions is still enigmatic.

How a given TF drives various transcriptional programs in different cell and tissue types is a longstanding question in Biology. It is assumed that *in vivo*, TFs establish dynamic protein-protein interaction (PPI) networks (Rhee et al., 2014). We tackled this issue using the Hox TFs, which are key players of animal development and cell homeostasis in adult. The Hox proteins belong to the conserved class of HD TFs (Bürglin and Affolter, 2016). They are expressed along the longitudinal axis in cnidarians (He et al., 2018) and bilaterians (Pearson et al., 2005), and orchestrate the development and homeostasis of a diversity of cell and tissue types (Castelli-Gair Hombría et al., 2016; Pearson et al., 2005). Understanding their molecular function has been a central aim in Developmental Biology. We previously used a tissue-specific proximity-labelling proteomic method to capture interactors of the Hox TF Ultrabithorax (Ubx) in *Drosophila* embryo (Carnesecchi et al., 2020). Our work revealed that the Ubx protein-networks are assembled at several layers of gene expression. Notably, many partners were regulators of mRNA splicing, revealing an unexpected aspect of the Hox operating mode *in vivo*. Based on our findings, we propose that Hox TFs coordinate gene expression programs by modulating transcription and splicing.

In this study, we combined transcriptomic profiling of differentially spliced genes in the embryonic mesoderm and in *Drosophila* S2R+ cells to uncover a novel function of Ubx in splicing. Notably, Ubx regulates mRNA expression and splicing to promote distinct functions in different cell and tissue contexts. Comparison of transcriptome and genome-wide chromatin binding profiles indicated that Ubx largely binds its spliced target genes in exons and introns. In line, transcriptome from *Drosophila* S2R+ cells expressing the Ubx^N51A^ mutant, which is no longer able to bind DNA, indicated that the Ubx HD is required for its splicing function. Furthermore, we uncovered a novel RNA-binding ability of Ubx in cells and *in vitro*. Intriguingly, the N51 amino acid of the HD is non-essential for mediating Ubx-RNA interface *in vitro* but is required for *in vivo* Ubx-RNA interaction. We found that the N51 amino acid mediates the interaction between Ubx and the active form (S2Phos) of Pol II on chromatin. By applying drugs or using Pol II mutant affecting the transcription rate, we uncovered a dynamic interplay between Ubx and Pol II that orchestrates elongation-coupled splicing. In sum, Ubx regulates splicing via its HD in a co-transcriptional manner. By extending the molecular repertoire of the Hox TFs, our work provides pivotal entry points to understand the Hox function in development and diseases, and unique view point on the role of TFs at the mRNA-regulatory level for governing cell fate decision.

## RESULTS

### Ubx modulates mRNA splicing *in vivo* to coordinate tissue-specific functions

We previously uncovered an interplay between the *Drosophila* Hox TF Ubx and splicing factors for muscle development (Carnesecchi et al., 2020). Based on these results we hypothesized that Ubx coordinates cell fate decision by modulating splicing.

To test this assumption, we analysed in depth our transcriptome data performed in differentiating mesoderm (stage 14-17) upon mesoderm-specific depletion of Ubx (Fig.1a) (Caussinus et al., 2011; Domsch et al., 2019). The transcriptome profile exhibited a high number of genes differentially expressed upon Ubx knock-down (KD) as previously described (Supplementary Fig.1a, Supplementary Table 1). These genes are referred to as transcriptionally regulated or misexpressed. We next asked whether mRNA splicing was affected by Ubx degradation. To this end, we analysed the transcriptomes with Junction-Seq (Hartley and Mullikin, 2016), a bioinformatic package that detects differential usage of exons and exon-exon junctions (splice sites). Moreover, Junction-Seq offers a visualisation of the transcript isoforms relative to the differential exon and junction usage, thereby providing information with regards to the splice events. We identified 425 differential splicing events upon Ubx degradation compared to control condition (Fig.1b-c, Supplementary Table 2). Among these events, 82% were differential exon usage (347/425) with 14% involving the first exon (49/347). The data exhibited a moderate difference of events for the exon (59% high, 41% low), as well as for the splice sites usages (40% high, 60% low, Fig.1c). Moreover, these events were related to a substantial number of genes differentially spliced upon Ubx degradation (Fig.1d, 283). We overlapped the list of genes differentially spliced and misexpressed upon Ubx depletion (Fig.1d). The data revealed that 70% of the genes (199/283) are uniquely regulated at the splicing level while 84 genes were differentially spliced and misexpressed upon Ubx degradation (Fig.1d).

**Figure 1:**
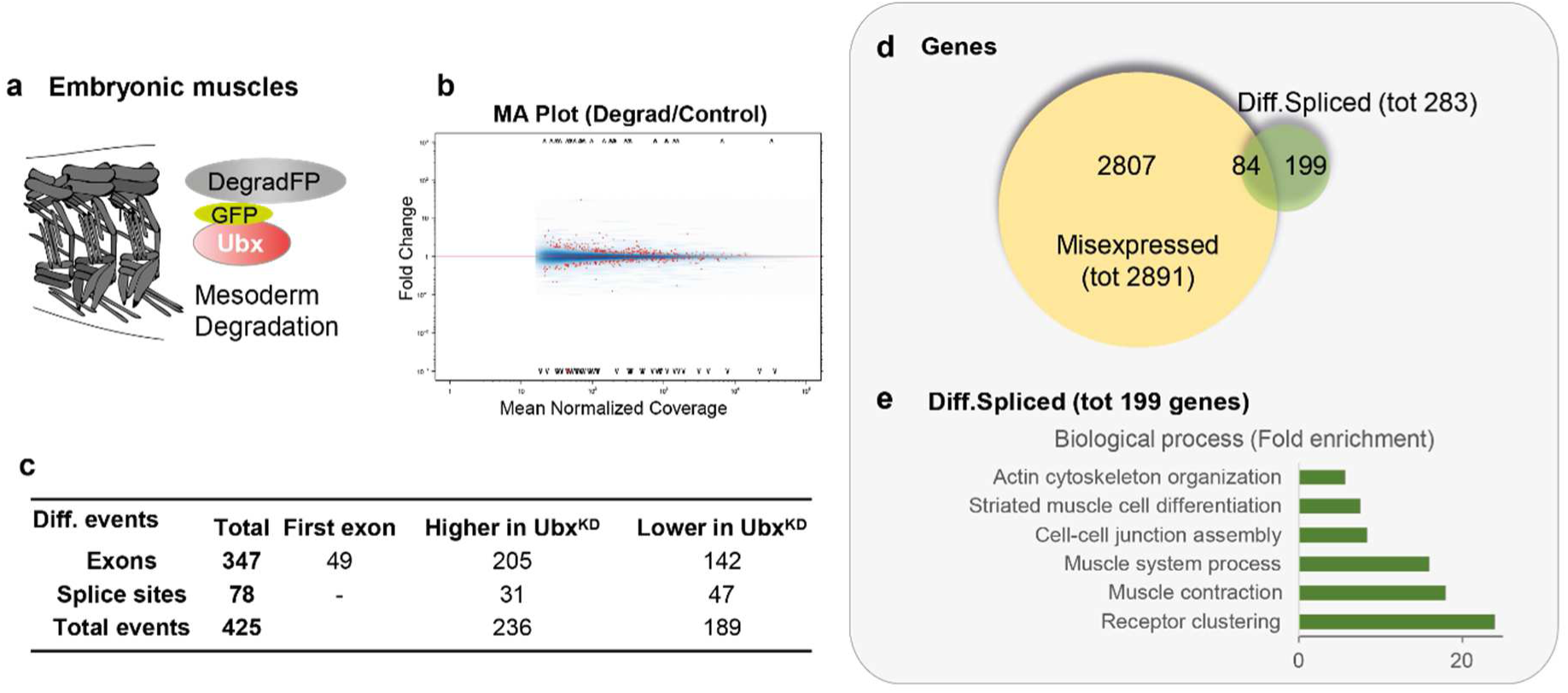
Ubx regulates splicing *in vivo* in the embryonic mesoderm. **a.** Schematic of tissue context for the differential transcriptome performed in embryonic mesoderm (Domsch et al., 2019), upon tissue-specific degradation of endogenous GFP-Ubx with the DegradFP system. **b.** MA plot from Junction-Seq shows the fold change of differential splicing events (higher/lower) in Ubx-degradFP (Degrad) or control experiment, plotted with the mean of the normalized coverage (FDR 0.1). **c.** Summary of differential splicing events detected upon Ubx degradation (exons, splice sites), higher or lower upon Ubx knock-down (KD). **d.** Venn diagram overlapping the misexpressed (2891) and differentially spliced (283) genes upon Ubx depletion in embryonic mesoderm. **e.** Fold enrichment of Gene Ontology (GO) term of biological processes enriched for the list of genes exclusively differentially spliced (199/283, pvalue<0.05, fold enrichment). See also supplementary Fig.1-3, Supplementary Table 1-2.

Focusing on the differentially spliced genes (199/283), Gene Ontology (GO) term analysis revealed an enrichment of biological functions related to striated muscle differentiation, muscle contraction, actin cytoskeleton and cell-cell junction assembly (Fig.1e). In contrast, the genes transcriptionally repressed by Ubx related to alternative cell fate, as previously described (Domsch et al., 2019) (Supplementary Fig.1b). The genes activated by Ubx (i.e., down-regulated upon Ubx degradation) were associated with general functions such as translation (Supplementary Fig.1c). This suggested that Ubx regulates mRNA expression and splicing to coordinate distinct cell functions. The molecular function (GO term) of the differentially spliced genes upon Ubx degradation were enriched for cytoskeletal proteins as well as Rho GTPase activity, which is essential for cell shape and motility (Ridley, 2001) (Supplementary Fig.1d). Consistently, cell components of the differentially spliced genes were related to neuromuscular junction, myofibril and sarcomere module (Supplementary Fig.1d). The list of genes included Tropomyosin I, Paramyosin, Troponin T as well as Complexin, Ephexin and Neto for neuromuscular junction (Supplementary Fig.2a-b). This strongly suggests that Ubx regulates muscle development not only by stabilising the lineage-specific transcriptional program but also by coordinating muscle function and signal transduction through the regulation of alternative splicing (AS) and specific isoforms production.

One of the various AS mechanisms relies on modulating the expression and/or splicing of mRNA regulatory proteins themselves (Smith and Valcárcel, 2000). In this context, we examined the genes related to mRNA processing, which were misexpressed and/or differentially spliced upon Ubx degradation in the mesoderm. The GO term analysis revealed a low enrichment of events related to mRNA processing among the genes repressed by Ubx (i.e., up-regulated upon Ubx degradation) (Supplementary Fig.1c). Moreover, Ubx depletion is associated with the differential splicing of a small set of mRNA processing factors, such as Squid (Sqd), a hnRNP for which isoform-specific functions have been described (Norvell et al., 1999), RbFox1, involved in cardiac hypertrophy and heart failure (Gao et al., 2015) or the serine threonine kinase DOA, producing multiple isoforms for which non-redundant functions have been suggested (Kpebe and Rabinow, 2008) (Supplementary Fig.3). Thus, Ubx could modulate splicing by regulating the level and splicing of mRNA processing factors, yet it does not exclude a direct effect of Ubx on splicing.

Overall, these data revealed that Ubx regulates tissue-specific functions through the regulation of mRNA expression and splicing. We had shown that Ubx regulates alternative fate gene expression at the transcriptional level (Domsch et al., 2019). Our results now suggest a novel regulatory function of Ubx in muscle development by coordinating gene programs at the splicing level.

### Ubx distinctly coordinates transcription and splicing

The mesoderm is composed of various cell lineages (somatic, visceral and cardiac). Importantly, the muscle types are specified by defined transcription and splicing programs (Nikonova et al., 2020; Spletter and Schnorrer, 2014). Consequently, the heterogeneity of the mesoderm population might shade the impact of Ubx in splicing. In addition, Ubx regulates the expression of mRNA processing factors thereby challenging the identification of its direct function in splicing. To decouple the role of Ubx on transcription and splicing, we chose to investigate its molecular function in a homogenous cell context, which does not express any Hox proteins. Thus, we investigated the role of Ubx in splicing using the Hox free S2R+ *Drosophila* cell system. Specifically, we performed RNA-Seq experiments upon ectopic expression of Ubx Wild-Type (WT) or GFP fused to a nuclear localisation sequence (nls) as control (Fig.2a). We transiently expressed the constructs under the control of the GAL4-UAS system (Brand and Perrimon, 1993) driven by the ubiquitous actin promoter. Three independent biological replicates were evaluated and Principle Component Analysis (PCA) validated the similarity of replicates (Supplementary Fig.4a). The differential expression profile revealed that ectopic expression of Ubx^WT^ induced a global change of the transcriptome compared to the control (GFP), with 932 up-regulated and 985 down-regulated genes (Supplementary Fig.4b-c, Supplementary Table 3). The transcriptome performed in the differentiating mesoderm revealed that Ubx represses alternative fate genes at the transcriptional level to stabilise the cell lineage (Domsch et al., 2019). In contrast, the *Drosophila* S2R+ cells represent a pool of somatic cells, which are not specified. In this context, the genes activated by Ubx^WT^ were largely related to various tissue identities and differentiation processes, such as heart development, motor neuron axon guidance, histoblast morphogenesis, stem cell differentiation. This mirrors the pivotal role of Ubx, and of the Hox TFs in general, in the development of numerous tissue types (Supplementary Fig.4d). It also demonstrates the inability of Ubx to control one cell-specific transcriptional program in absence of the proper set of TFs essential for lineage commitment (Seifert, 2015). In contrast, Ubx^WT^ repressed genes related to general processes such as translation and ribosome biogenesis in this cellular context (Supplementary Fig.4e). Subsequent analysis using Junction-Seq revealed that Ubx^WT^ expression induced a significant change of the mRNA splicing profile of the cells, with 133 events differentially regulated compared to control (Fig.2b-c, Supplementary Table 4). Similar to the mesoderm transcriptome (Fig.1), we observed 85.7% of differential exons usage (114/133) within 19% involving the first exon (Fig.2c, 22/114). The data indicated a moderate difference of events upon Ubx expression for the splice sites usage (58% high, 42% low), however the difference was noticeable for the exon usage, with 68% of events higher upon Ubx^WT^ expression (Fig.2c). This strongly suggests that Ubx largely promotes the retention of exon cassettes in *Drosophila* cells. These events related to 81 genes differentially spliced upon Ubx^WT^ expression (Fig.2d). We confirmed the role of Ubx on splicing by analysing the differential exon retention of several target genes upon Ubx^WT^ expression, namely Chascon (Chas, Fig.2f, i-j), the poly(A) binding protein pAbp (Fig.2g, k-l), the small GTPase Rgk1 (Fig.2h, m-n), the RhoGEF Puratrophin-1-like (Pura, Supplementary Fig.5a, c-d), the histone H3.3B (Supplementary Fig.5b, e-f), the cAMP phosphodiesterase Dunce (Dnc, Supplementary Fig.5g-h), CG34417 (Supplementary Fig.5i-j) and the ribosomal protein Rps13 (Supplementary Fig. 5k-l). This is exemplified with Chas, a gene involved in hair bristle development and muscle-tendon junction (Olguín et al., 2011), which is regulated by Ubx, both at the transcriptional and splicing level (Fig.2f, i-j). Chas contains an exon cassette retained upon Ubx^WT^ expression (Fig.2f, exon E5). We confirmed the result by assessing the expression level of constitutive and alternative exons related to the housekeeping gene Actin5C. This showed a change at the expression level for all exons tested (E1, E3, E5, Fig.2i). In contrast, pAbp was regulated at the splicing level exclusively, and only exon cassette E1 was significantly modulated at the expression level (Fig.2k). We further examined the differential splicing of the exon cassette E5 of Chas compared to its constitutive exon E3 showing an increase of exon E5 retention (Fig.2j). In contrast, the constitutive exon E1 of Chas was not differentially spliced upon Ubx^WT^ expression.

**Figure 2:**
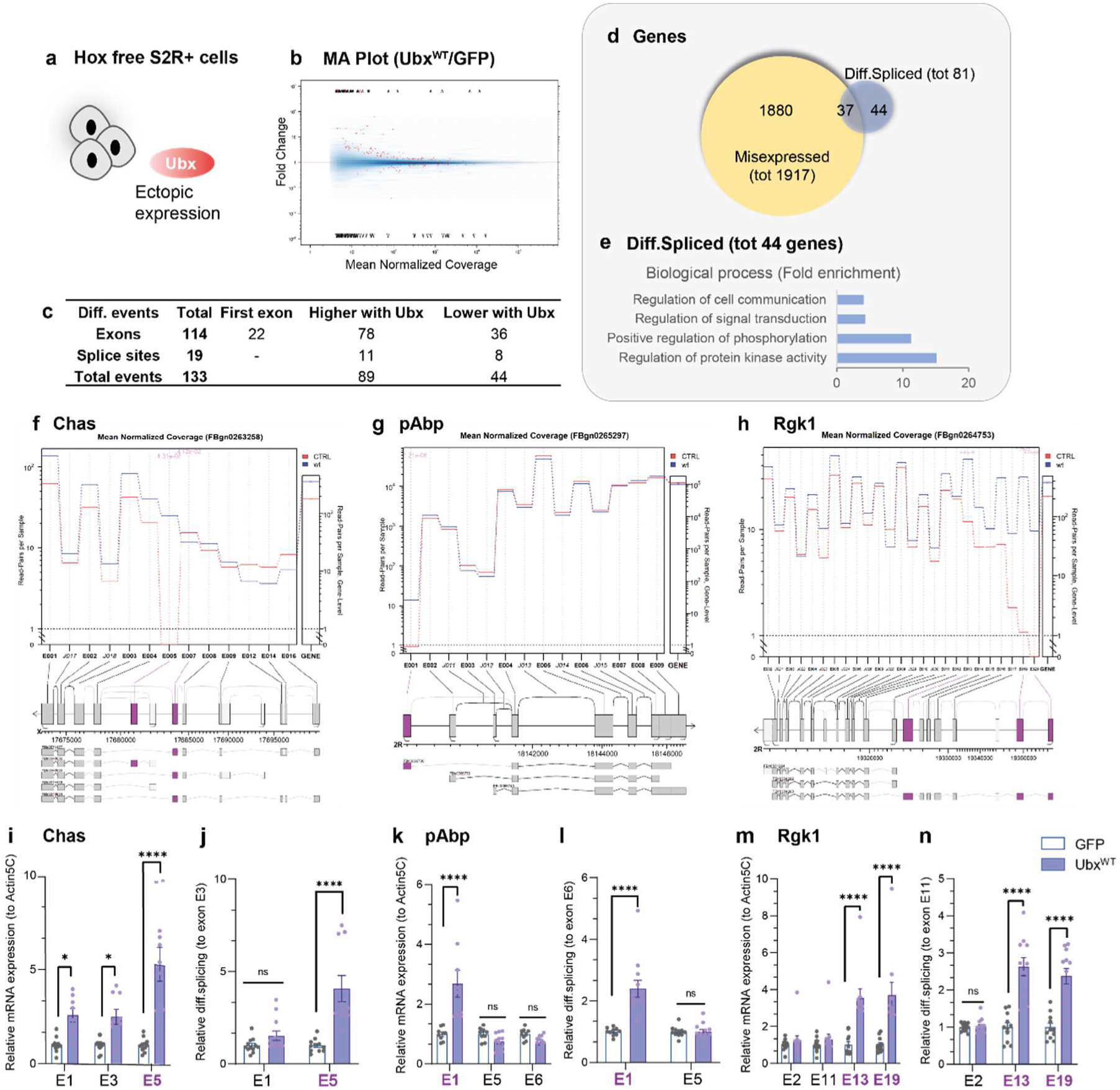
Ubx regulates transcription and splicing in S2R+ *Drosophila* cells. **a.** Schematic of cell context for the differential transcriptome performed in Hox free S2R+ *Drosophila* cells, upon ectopic expression of myc-Ubx^WT^ or GFP fused to a nuclear localisation sequence (nls) control, using the Gal4-UAS system driven by the actin promoter. RNA-Seq data were further analysed for 3 independent biological replicates. **b.** MA plot from Junction-Seq showed the fold change of differential splicing events (higher/lower) upon ectopic expression of Ubx^WT^ or GFP control in *Drosophila* S2R+ cells, plotted with the mean of the normalized coverage (FDR 0.1). **c.** Summary of differential splicing events detected upon Ubx^WT^ expression (exons, splice sites), higher or lower compared to control. **d.** Venn diagram overlapping the misexpressed (1917) and differentially spliced (81) genes upon Ubx^WT^ expression in *Drosophila* S2R+ cells. **e.** Gene ontology (GO) term of biological processes enriched for the list of genes exclusively differentially spliced (44/81, pvalue<0.05, fold enrichment). **f-h.** Visualisation from Junction-Seq of the mean normalised read count for each exon or splice junction (left Y-axis, extended panel), and the gene level normalised read count (right Y-axis, narrow panel) of Chas (**f**), pAbp (**g**) and Rgk1 (**h**), differentially spliced upon Ubx^WT^ expression (wt, blue line) compared to control (CTRL, red line). Significant differential splicing events are highlighted in purple. Isoforms including differentially spliced exon or junction usages (purple) upon Ubx^WT^ expression are displayed below each read count. **i-n**. RT-qPCR experiments showing the differential expression (**i, k, m**) over actin5C and **(j, l, n**) differential retention of exon cassettes over constitutive exons for Chas (**i-j**), pAbp (**k-l**) and Rgk1 (**m-n**) in *Drosophila* S2R+ cells expressing GFP control or Ubx^WT^ (blue). (E+number)=exon related to Junction-Seq annotation. Differentially spliced exons are underlined (purple). n=4 independent biological triplicates. Bars represent mean±SEM. Statistical test by one-way ANOVA (p<0.05 *, p<0.01**, p<0.001***, p<0.0001****, ns=non-significant. See also supplementary Fig.4-5, Supplementary Table 3-4.

We next overlapped the lists of genes misexpressed and differentially spliced upon Ubx^WT^ expression (Fig.2d). 54% of the genes (44/81) were exclusively differentially spliced without their expression being affected, indicating that Ubx regulates both transcription and splicing of mRNA. Interestingly, these genes related to cell function similar to the genes differentially spliced in the mesoderm, notably for cell communication, signal transduction and cell migration (Fig.2e, Supplementary Fig.4f). As observed for the mesoderm transcriptome, these functions were distinct from the one related to the misexpressed genes (Supplementary Fig.4d-e). This indicated once more that Ubx coordinates transcription and splicing to control distinct cell functions in defined cell and tissue types. Notably, its regulatory role in splicing is essential for cell-cell communication and cell shape. Consistently, we identified 16 genes differentially spliced both in the embryonic mesoderm and *Drosophila* S2R+ cells, highlighting common as well as cell-type specific target genes related to specialised cell behaviour (Supplementary Fig.4g).

Taken together, these results showed that Ubx controls transcription and splicing to regulate specific set of cellular functions in precise cell and tissue contexts.

### Ubx chromatin-binding events are enriched in the gene body of target genes

Ubx regulates its direct transcriptional targets by specific recognition and binding of cis-regulatory elements (Agrawal et al., 2011; Domsch et al., 2019; Mann and Chan, 1996). We thus reasoned that Ubx could modulate mRNA splicing through direct binding of its target genes. To study this question, Ubx chromatin binding profiles performed in *Drosophila* S2 cells (ChIP-Seq, Zouaz et al., 2017, Fig.3a) were compared to the transcriptome and splicing profiles upon Ubx^WT^ expression. The overlap showed that 71.5% (1372/1917) of the misexpressed and 85% (69/81) of the differentially spliced genes upon Ubx expression were bound by Ubx (Fig.3b). This indicated that these genes are most likely direct targets of Ubx. Previous analysis showed that Ubx binding is enriched at promoters (Zouaz et al., 2017). Consistently, we detected 41% of the binding events at promoters (promoter and Transcription Start Site TSS) (Fig.3a). Importantly, Ubx binding events were highly enriched within the gene body (48%), in particular in introns (73%), at Transcription Termination Sites (TTS 16%) and in exons (10%). In this context, we asked whether Ubx could be enriched in the gene body of its differentially spliced target genes (Fig.3b-c). The genomic distribution analysis revealed that the genes differentially spliced had an enrichment of binding events in introns (21/29%) and exons (8/16%) compared to all genes bound (Fig.3c, Chi^2^ p=0.0017). This difference was moderate compared to the genes misexpressed upon Ubx^WT^ expression (Chi^2^ p=0.1). Remarkably, Ubx binding events were not enriched in alternative exon cassettes (6/24 total exons). Instead, we noticed a broad distribution of Ubx along the gene body of various target genes, as exemplified for Chas, Rgk1 and Pura (Fig.3d, Supplementary Fig.6a-c).

**Figure 3:**
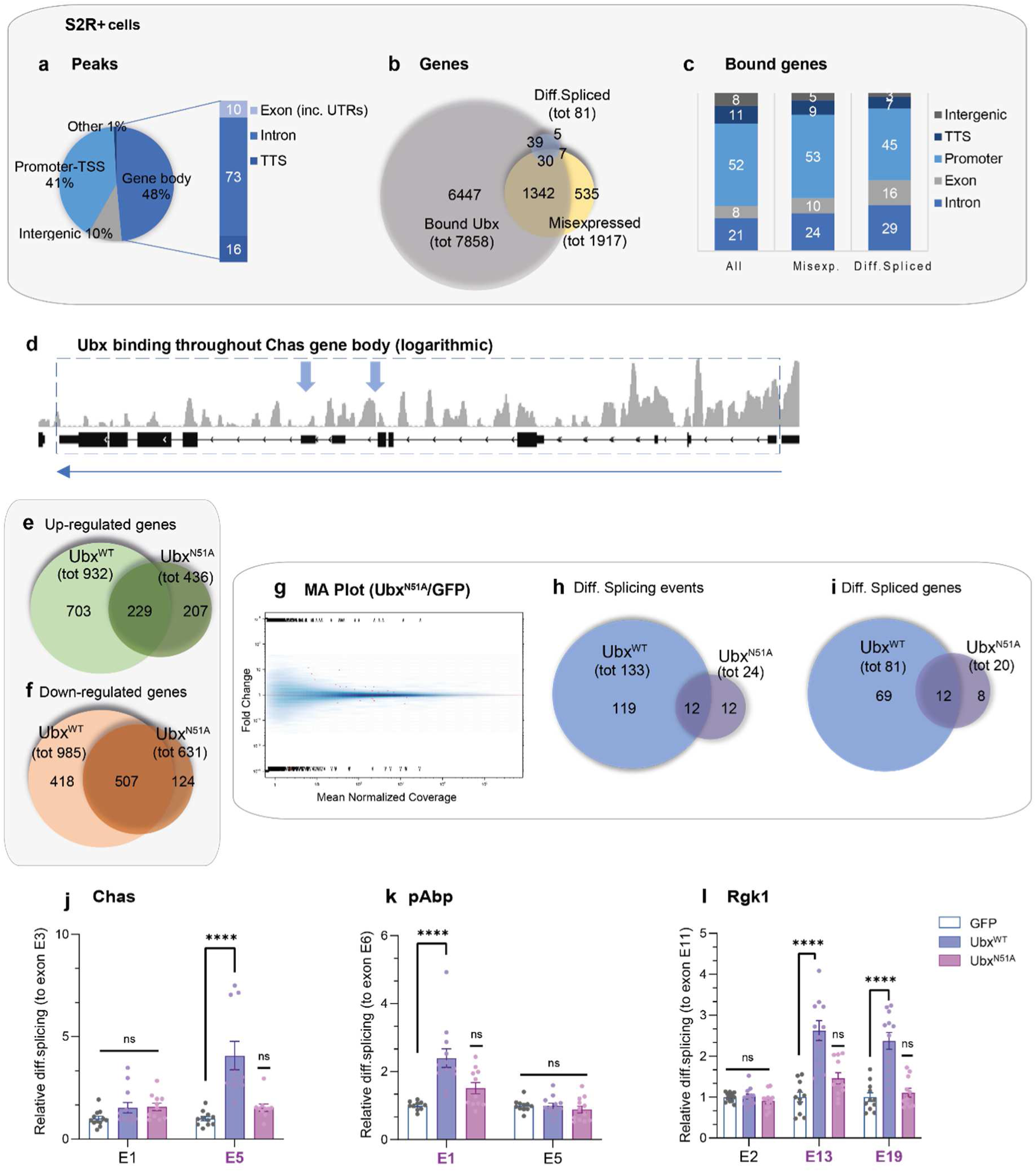
Ubx regulates splicing through its DNA-binding ability. **a.** Distribution of the genomic regions bound by Ubx, namely promoter and TSS (Transcription Start Site), intergenic, gene body, using the ChIP-Seq data of Ubx from *Drosophila* S2 cells generated by Zouaz et al. (2017). The distribution in the gene body is further detailed for intron, Transcription Termination Sites (TTS) and exons (including 5’ and 3’ UTRs). **b.** Venn diagram representing the overlap of genes bound, misexpressed and differentially spliced upon Ubx^WT^ expression in *Drosophila* cells. **c.** Graphical view of the distribution (%) of the total genes bound, misexpressed and bound, differentially spliced and bound by Ubx^WT^ according to intergenic, promoter, exon, intron and TTS. Chi^2^ tests were performed to test the distribution: “all vs misexp.”: p=0.596, “all vs diff.spliced”: p=0.0017, “misexp. vs diff.spliced”: p=0.12. **d.** Visualisation of Ubx binding events in the gene body of Chas by logarithmic scale. Transcription directionality is indicated with blue arrow. **e-f.** Venn diagrams from the RNA-Seq data performed in *Drosophila* S2R+ cells, representing the overlap of genes up-regulated (**e**) and down-regulated (**f**) upon Ubx^WT^ and Ubx^N51A^ expression compared to control GFP (3 independent biological replicates for each sample). **g.** MA plot from Junction-Seq showed the fold change of differential splicing events (higher/lower) upon ectopic expression of Ubx^N51A^ or GFP control in *Drosophila* S2R+ cells, plotted with the mean of the normalised coverage (FDR 0.1). **h-i.** Venn diagram representing the overlap of the differentially spliced events (**h**) and differentially spliced genes upon Ubx^WT^ and Ubx^N51A^ expression in *Drosophila* S2R+ cells. **j-l.** Relative RNA expression (RT-qPCR) revealed the differential retention of exon cassettes over constitutive exons for Chas (**j**), pAbp (**k**) and Rgk1 (**l**) in *Drosophila* S2R+ cells expressing GFP control, Ubx^WT^ (blue), or Ubx^N51A^ (purple), but not for constitutive exons. (E+number)=exon relates to Junction-Seq annotation, differentially spliced exons are underlined (purple). n=4 independent biological triplicates. Bars represent mean±SEM. Statistical test by one-way ANOVA (p<0.0001****, ns=non-significant). See also supplementary Fig.6-8, Supplementary Table 5-6.

Similarly, the analysis of Ubx binding profile in the mesoderm (stage 14-17) revealed that 56% of the binding events were localised in the gene body, including 60% in introns, 16% at TTS and 23% in exons (Supplementary Fig.6d). Notably, 50% of the differentially spliced genes upon Ubx degradation had an Ubx binding event (Supplementary Fig.6e). Moreover, we observed an enrichment of Ubx events in the gene body of the genes differentially spliced compared to the genes only bound or regulated at the transcriptional level (Chi^2^ p=0.0005, p=0.08, Supplementary Fig.6f). Although the distribution of events was unchanged in exons, more events were identified in introns (26/42%) and less in intergenic regions (18/9%). This indicated an enrichment of Ubx binding events within the gene body (Supplementary Fig.6f).

Overall, the analysis strongly suggested that Ubx regulates transcription as well as splicing via direct chromatin binding of its target genes. Notably, Ubx binding is enriched along the gene body of its spliced target genes.

### Ubx splicing activity requires the Homeodomain

The Hox DNA-binding domain, the Homeodomain (HD), plays a pivotal role in the DNA recognition, binding and regulatory function of Hox TFs (Berger et al., 2008; Mann and Chan, 1996). Thus, the HD could be an essential platform for Ubx transcriptional and splicing functions. To evaluate the role of Ubx binding ability in splicing, we performed transcriptome profiling upon transient expression of Ubx^N51A^, a mutant version of Ubx which is no longer able to bind DNA (Carnesecchi et al., 2020). N51A (asparagine to alanine) is a single mutation of the amino-acid N51 of the HD that is necessary for the interaction with the major groove of DNA (Passner et al., 1999). The Ubx^N51A^ mutant has been generated and validated in our previous study, showing its inability to induce homeotic transformation in embryos or activate Ubx synthetic enhancer in S2R+ *Drosophila* cells (Carnesecchi et al., 2020).

RNA-Seq experiments were performed in *Drosophila* S2R+ cells expressing Ubx^N51A^ and PCA component analysis validated the high similarity of the three biological replicates (Supplementary Fig.7a). The differential expression profile revealed that ectopic expression of Ubx^N51A^ induced a global change of gene expression compared to control, with 436 up-regulated and 631 down-regulated genes (Supplementary Fig.7b, Fig.3e-f, Supplementary Table 5). Intriguingly, 22.8% (213/932) of the up-regulated genes and 50.2% (495/985) of the down-regulated genes upon Ubx^WT^ expression were equally regulated by Ubx^N51A^ mutant (Fig.3e-f). This could be due to an indirect effect or residual chromatin loading via protein-protein interactions (Beh et al., 2016).

Analysis using Junction-Seq revealed that Ubx^N51A^ expression induced rare differential splicing events with only 24 events detected (Fig.3g-h, Supplementary Fig.7c), in 20 genes (Fig.3i, Supplementary Table 6). This demonstrated that Ubx^N51A^ has a minor effect on splicing. Comparison of differential splicing profiles revealed that 89% (119/133) of the splicing events and 85% (69/81) of differentially spliced genes were exclusively regulated by Ubx^WT^ (Fig.3h-i). This difference did not account for Ubx expression level as both Ubx^WT^ and Ubx^N51A^ constructs were expressed at comparable level (Supplementary Fig.7d). We confirmed this finding by analysing the differential exon retention of selected target genes upon Ubx^N51A^ expression (Fig.3j-l, Supplementary Fig.8). As expected, Ubx^N51A^ did not promote the differential splicing of Chas, Rgk1, pAbp, Pura and H3.3B. Moreover, Dnc, one of the common spliced target genes of Ubx^WT^ and Ubx^N51A^ exhibited a differential retention of exon E11 upon expression of each Ubx form (Supplementary Fig.8h-i). Interestingly, some of the genes repressed both by Ubx^WT^ and Ubx^N51A^ are splicing factors (Rm62, SF3B3). It indicated once more that while Ubx regulates the expression of splicing factors, its activity on alternative splicing is most likely mediated by direct effect on mRNA targets.

Taken together, the data showed that Ubx splicing activity required a functional homeodomain, while its transcriptional activity was partly mediated by the Ubx^N51A^. This strongly suggests that the full binding ability of Ubx HD is essential for its splicing regulatory function.

### Ubx binds RNA *in vivo* and is enriched on target alternative exon cassettes

Our data indicated that N51 of the HD is necessary for Ubx splicing activity, yet we did not observe a specific enrichment of Ubx DNA-binding events on specific exon cassette (Fig.3d, Supplementary Fig.6a-c). Therefore, we wondered if the splicing activity of Ubx could be mediated by a so far uncovered RNA-binding ability thereby providing specificity at the exon level.

To this end, we performed RNA-ImmunoPrecipitation experiments of GFP fused proteins (RIP-RTqPCR) using *Drosophila* cells nuclear extracts. We reasoned that nuclear extracts exhibit the RNA-binding function linked to transcription and mRNA processing, while interactions occurring in the cytoplasm are most likely related to mRNA transport and translation. We assessed the enrichment of constitutive and alternative exons in GFP, GFP- Ubx^WT^ and GFP-Ubx^N51A^ fractions for Ubx spliced genes Chas, pAbp, Rgk1 (Fig.4a-c) and Pura, H3.3B (Supplementary Fig.9a-b). The enrichment of Actin5C mRNA was measured as a negative control (Fig.4a). For all target genes, we observed an enrichment of exon cassettes (Chas E5, pAbp E1, Rgk1 E13-E19, Pura E11, H3.3B E4) in the Ubx^WT^ fraction compared to control GFP. This result exhibits a novel ability of Ubx to interact with RNA *in vivo*. Interestingly, the constitutive exons of Chas, pAbp, Rgk1 and Pura were weakly or to a lesser extent enriched in Ubx^WT^ fraction compared to exon cassettes (Fig.4a-c, Supplementary Fig.9a). This indicated a binding specificity of Ubx toward alternatively spliced exons. Only H3.3B presented a similar enrichment for both constitutive E2 and cassette E4 exons in Ubx^WT^ fraction (Supplementary Fig.9b). The smaller mRNA size of H3.3B compared to the other mRNA targets (mRNA H3.3B=1.3kb while other mRNAs>2.4kb) could account for this observation. Importantly, we observed a significant decrease of RNA pull-down in the Ubx^N51A^ fraction compared to Ubx^WT^ for all RNA targets studied (Fig.4a-c, Supplementary Fig.9a-b). As control, we analysed the pull-down efficiency by immunoblotting, showing a similar enrichment of GFP-fused proteins (Supplementary Fig.9c).

**Figure 4:**
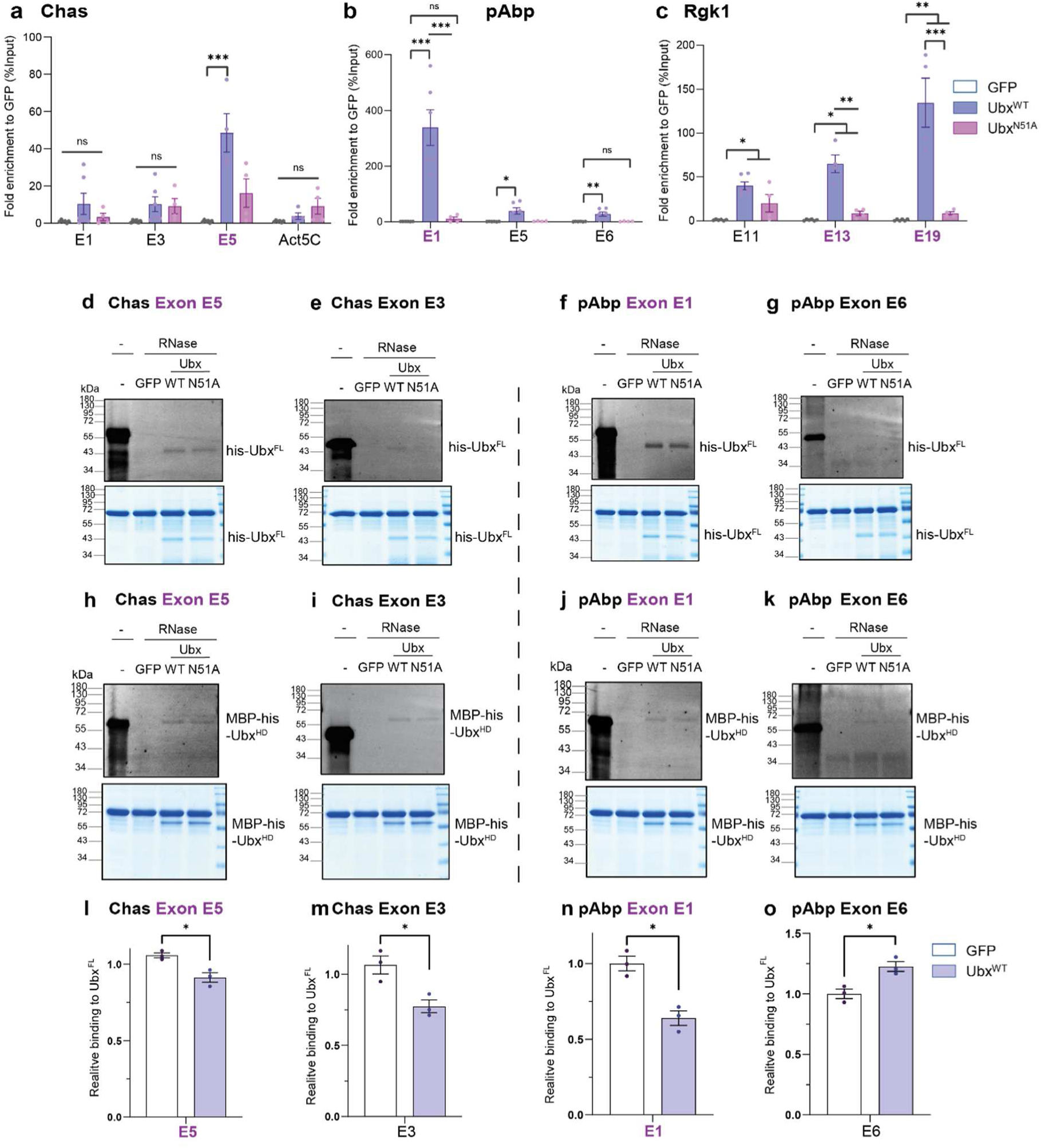
Ubx binds RNA *in vivo* and *in vitro* and N51 of the homeodomain is non-essential for its RNA-binding abilities. **a.** RNA-Immunoprecipitation (RIP-RTqPCR) experiments of *Drosophila* S2R+ cells expressing GFP control, GFP-Ubx^WT^ (blue) and GFP-Ubx^N51A^ (purple) showing an enrichment of targeted exonic regions of Chas (**a**), pAbp (**b**) and Rgk1 (**c**). Values are RNA relative enrichment over GFP calculated as percentage of input. (E+number)=exon related to Junction-Seq annotation, differentially spliced exons are underlined (purple). n=3 independent biological duplicates. Bars represent mean±SEM. The results exhibited a specific enrichment of differentially spliced exonic sequences in Ubx^WT^ fraction compared to GFP and Ubx^N51A^. **d-k.** Fluorescent protein-RNA interaction assay followed by UV-crosslinking and RNase digestion, performed *in vitro* with purified proteins namely, MBP-his-GFP as control, his-Ubx (WT and N51A) full-length (FL) (**d-g**) and the homeodomain alone MBP-Ubx-HD (WT and N51A) (**h-k**). Interactions were detected on denaturating gels by Cy3-UTP signal (upper panel), and gels were stained by Coomassie to reveal the protein content (lower panel). Each probe is indicated relative to the genes and exons. Molecular marker is indicated. **l-o.** Graphical view showing the quantification of relative RNA-binding of Ubx homeodomain (HD) compared to full-length (FL) protein for Chas exon cassette E5 (**l**), constitutive E3, (**m**), pAbp exon cassette E1 (**n**), constitutive E6 (**o**) normalized to Coomassie staining. n=3 independent biological replicates. Statistical test by one-way ANOVA (p<0.05 *, p<0.01**, p<0.001***, ns=non-significant). See also supplementary Fig.9-10, Supplementary Table 7.

In sum, the results indicated that Ubx binds RNA and that the N51A mutation impacts on its RNA-binding ability *in vivo*. More importantly, the results showed that Ubx is specifically enriched in mRNA exonic regions regulated at the splicing regulatory level by Ubx, which contrasts with its DNA-binding profiles spread over the gene body.

### Ubx binds RNA directly *in vitro* and HD-N51 is a non-essential amino acid

We subsequently asked whether Ubx could directly bind RNA. To this end, we performed *in vitro* protein-RNA interaction assays by Ultra Violet (UV)-crosslinking using purified his-tagged proteins and fluorescent labelled RNA probes. The probes corresponded to the exact or nearest exonic sequences (if UTP-content was too low) identified in cells, and were labelled using Cy3-UTP nucleotides (Supplementary Table 7, 80-157 nucleotides length). Notably, we chose RNA sequences that contain similar UTP content, with a broad distribution of the nucleotide along the RNA probes (Supplementary Table 7). After crosslinking of protein-RNA complexes, unprotected RNAs were digested with RNase, and protected bound RNAs were visualised on denaturing gels for fluorescent RNA detection and protein content by Coomassie staining (Fig.4d-k, Supplementary Fig.9-10). Remarkably, the assays revealed a direct binding of purified Ubx protein on the RNA probes for the exon cassettes of Chas (E5), pAbp (E1), Rgk1 (E19) and Pura (E11) compared to GFP control (Fig4.d, f, Supplementary Fig.9d, f). In contrast, Ubx binding was weaker for the probes of constitutive exon for Chas (E3), Rgk1 (E1), Pura (E19) (Fig.4e, Supplementary Fig.9e, g) and not detected for pAbp (E6) (Fig.4g). Although we cannot account for protein-RNA interaction in UTP-free sequences, this suggested that Ubx exhibits binding specificity toward RNA sequences.

Concurrently, we analysed the RNA-binding ability of Ubx^N51A^ mutant. Surprisingly, Ubx^N51A^ exhibited similar RNA-binding ability as Ubx^WT^, which was supported by the quantification of protein-RNA interactions (Fig.4d-k, Supplementary Fig.9d-k, m-p). In contrast to the Ubx-RNA binding profile in *Drosophila* cells, the *in vitro* interaction assay revealed that the N51 amino-acid is non-essential for Ubx RNA-binding in sharp contrast with its DNA-binding ability (Supplementary Fig.9f). To evaluate if the HD domain mediates the interaction, we performed UV-crosslinking assay with purified Ubx-HD (Fig.4h-k, Supplementary Fig.9h-k). The assays revealed that Ubx HD^WT^ and HD^N51A^ both bound RNA. Further quantifications revealed that the Ubx HD did not recapitulate the full binding of the full-length protein for Chas exons E5 and E3, pAbp exon E1 (Fig.4l-n, Supplementary Fig.10a-c), Rgk1 exons E19, E1 and Pura exons E11, E19 (Supplementary Fig.10e-l). Unexpectedly, the HD alone provided a clear binding affinity of Ubx toward the RNA probes of pAbp constitutive exon E6 which was not detected with the full-length protein (Fig.4o, Supplementary Fig.10d). This indicated that while the HD of Ubx mediates RNA-binding, it is not sufficient to recapitulate the RNA-binding profile of the full-length Ubx protein.

In sum, these results revealed a novel ability of Ubx to directly bind RNA *in vitro*. Importantly, the N51 amino-acid of the HD is not essential for its direct RNA-binding. This strongly suggests that Ubx recognises and binds RNA and DNA via its homeodomain yet, using a different protein-RNA interaction mode.

### Ubx interacts with active RNA Polymerase II and requires its functional HD

Our results indicated that the N51 amino-acid of the HD is necessary for Ubx RNA-binding and splicing function *in vivo,* but is not essential for its RNA interaction *in vitro*. It suggests that Ubx loading onto the chromatin mediates its RNA interaction *in vivo* and regulates splicing. In this context, we explored in depth the molecular mechanism by which Ubx controls splicing.

Splicing happens mainly co-transcriptionally and depends on the RNA Polymerase II (Pol II) activity (Bentley, 2014). Moreover, Ubx interacts with several components of the basal transcriptional machinery (Baëza et al., 2015; Boube et al., 2014) and can modulate transcription events, such as Pol II pausing (Zouaz et al., 2017). Based on these evidences, we reasoned that Ubx and Pol II could collaborate for regulating splicing during active elongation process. Co-immunoprecipitation experiments were performed on GFP-fused proteins in *Drosophila* S2R+ cells, revealing that Ubx^WT^ interacts with both paused (initiation, S5Phos) and active (elongation, S2Phos) Pol II (Fig.5a-b). In contrast, Ubx^N51A^ mutant interacted similarly with the paused Pol II (S5Phos), while exhibiting a weak interaction with active Pol II (S2Phos) (Fig.5c). Notably, co-immunoprecipitation performed on embryos nuclear extract revealed that Ubx interacts equally well with the two phosphorylated forms of Pol II in embryos (Supplementary Fig.11).

**Figure 5:**
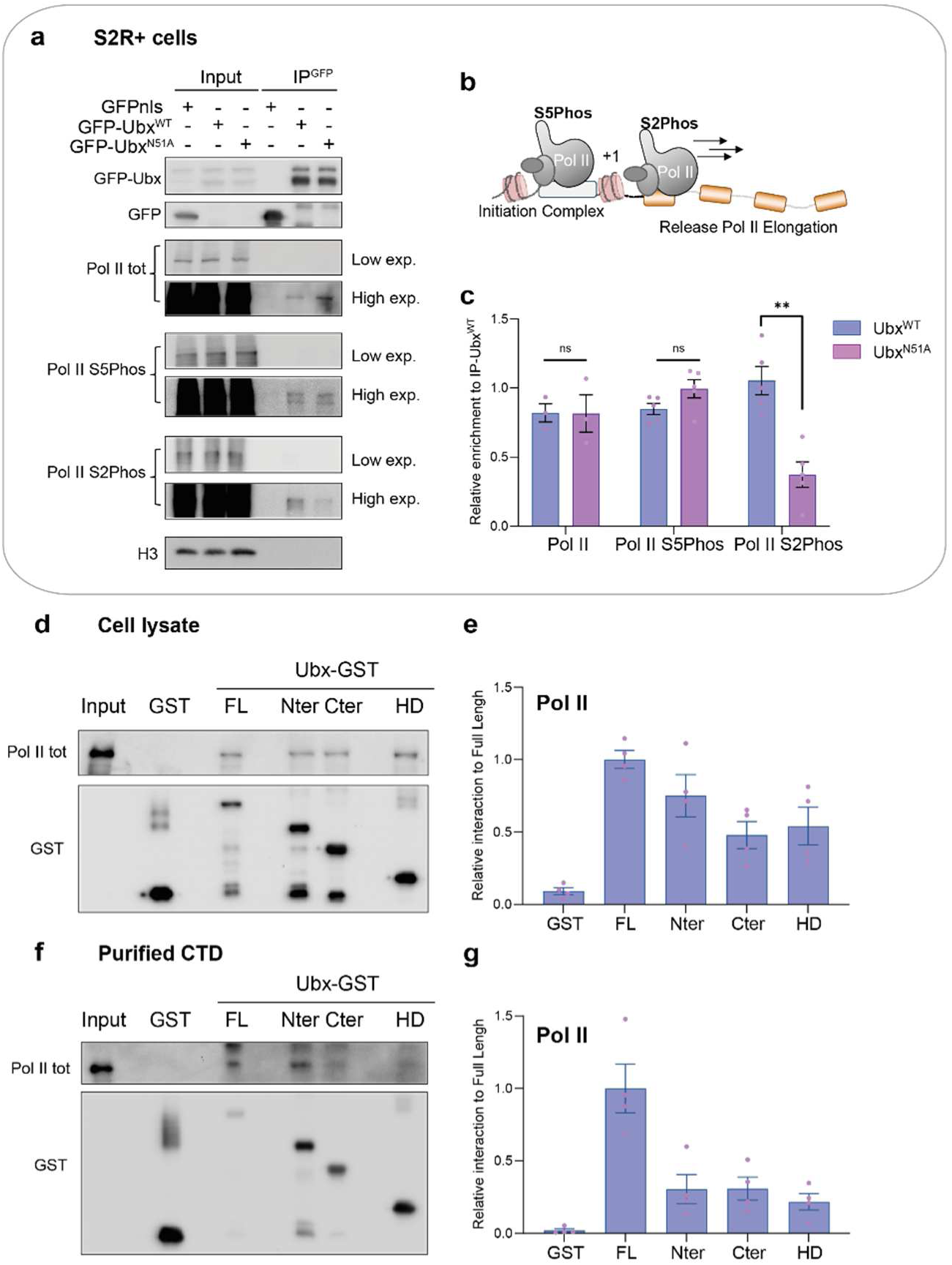
Ubx interacts with active RNA Polymerase II via its functional Homeodomain. **a.** Co-immunoprecipitation of endogenous Pol II total, paused (S5Phos) and active (elongating, S2Phos) forms with GFP fusion proteins (GFPnls, GFP-Ubx^WT^, GFP-Ubx^N51A^), ectopically expressed in *Drosophila* S2R+ cells. Western blots were probed with the indicated antibodies. The input is shown as a control of expression levels (lanes 1-3), Histone 3 (H3) is used as a loading control. Low and high exposure (exp.) are presented. **b.** Schematic of paused Pol II (S5phos) in the initiation complex, loaded onto the promoter and elongating Pol II (S2Phos) actively transcribing the gene. **c.** Quantification of relative enrichment of Pol II, Pol II S5phos and Pol II S2Phos relative to Ubx^WT^ and Ubx^N51A^ pulled down proteins, showing a specific enrichment of active Pol II S2Phos for Ubx^WT^ compared to Ubx^N51A^. n=3 independent biological replicates. Statistical test by one-way ANOVA (p<0.01**, ns=non-significant). **d-g.** Pull-down assay using the indicated GST-fused Ubx derivatives (Full-Length FL, Nter/N-terminal, C-ter/C-terminal, Homeodomain HD) and *Drosophila* S2R+ nuclear extracts (**d**) or *in vitro* purified human Carboxy Terminal Domain (CTD) of Pol II (**f**). Input is loaded as indicated. **e-g.** Quantification of interactions relative to GST-Ubx^FL^ (full-length) signal is indicated in (**e**) for Pol II from nuclear extract and in (**g**) for purified CTD. n=3 independent biological replicates. Bars represent mean±SEM. See also supplementary Fig.11.

We subsequently asked whether Ubx interaction with Pol II required a specific domain. To this end, GST-pull down experiments were performed using Ubx full-length (FL), truncated N-terminal, C-terminal or homeodomain (HD) purified proteins with *Drosophila* S2R+ cell extracts (Fig.5d-e). The assay revealed that Ubx full-length protein efficiently pulled down Pol II from nuclear extract, while each fragment pulled down Pol II to a lesser extent (50-80% of the full-length interaction). Interestingly, we noticed a stronger interaction of Pol II with the N-terminal domain of Ubx compared to the HD-containing C-terminal domain (80/50% Fig.5e). We subsequently examined if Ubx could directly interact with Pol II, a so far unknown molecular feature of the Hox TFs. We performed GST-pull down assay with Ubx derivatives and the purified human Carboxy Terminal Domain (CTD) of Pol II (Fig.5f-g). We observed a direct interaction between the CTD of Pol II and Ubx full-length protein as well as for the N-terminal, C-terminal and HD domain. The pull-down was ten times stronger with Ubx full-length than with the Ubx truncated fragments (Fig.5g). We noticed a stronger interaction with the N-terminal domain compared to the C-terminal, however, not significantly distinguished after quantification (n=4).

In sum, the data showed that Ubx interacts with Pol II *in vitro* and *in vivo* and that the N51 amino-acid is essential for the interplay between Ubx and active Pol II. This strongly suggests that Ubx binding to the chromatin is essential to mediate Ubx/Pol II interaction during active transcription.

### Ubx regulates splicing via elongation-mediated process

We next asked whether the Pol II activity can impact on Ubx splicing function. To do so, we first assessed the interaction between Ubx and Pol II upon treatment with the transcription inhibitor Actinomycin D, a DNA intercalator that accumulates hyperphosphorylated Pol II (Fig.6a-d (Bensaude, 2011)). Actinomycin D treatment reduced the interactions between Ubx^WT^ or Ubx^N51A^ with paused Pol II (S5Phos, 20%), and even more between Ubx^WT^ and active Pol II for which the interaction was reduced by 70% (Fig.6d). As a control, the input fraction confirmed that this effect was not due to a decrease of Pol II phosphorylation (Fig.6a). This indicated that Ubx/Pol II interaction depends on active transcription as well as on the DNA-binding ability of Ubx as previously highlighted (Fig.6d).

**Figure 6:**
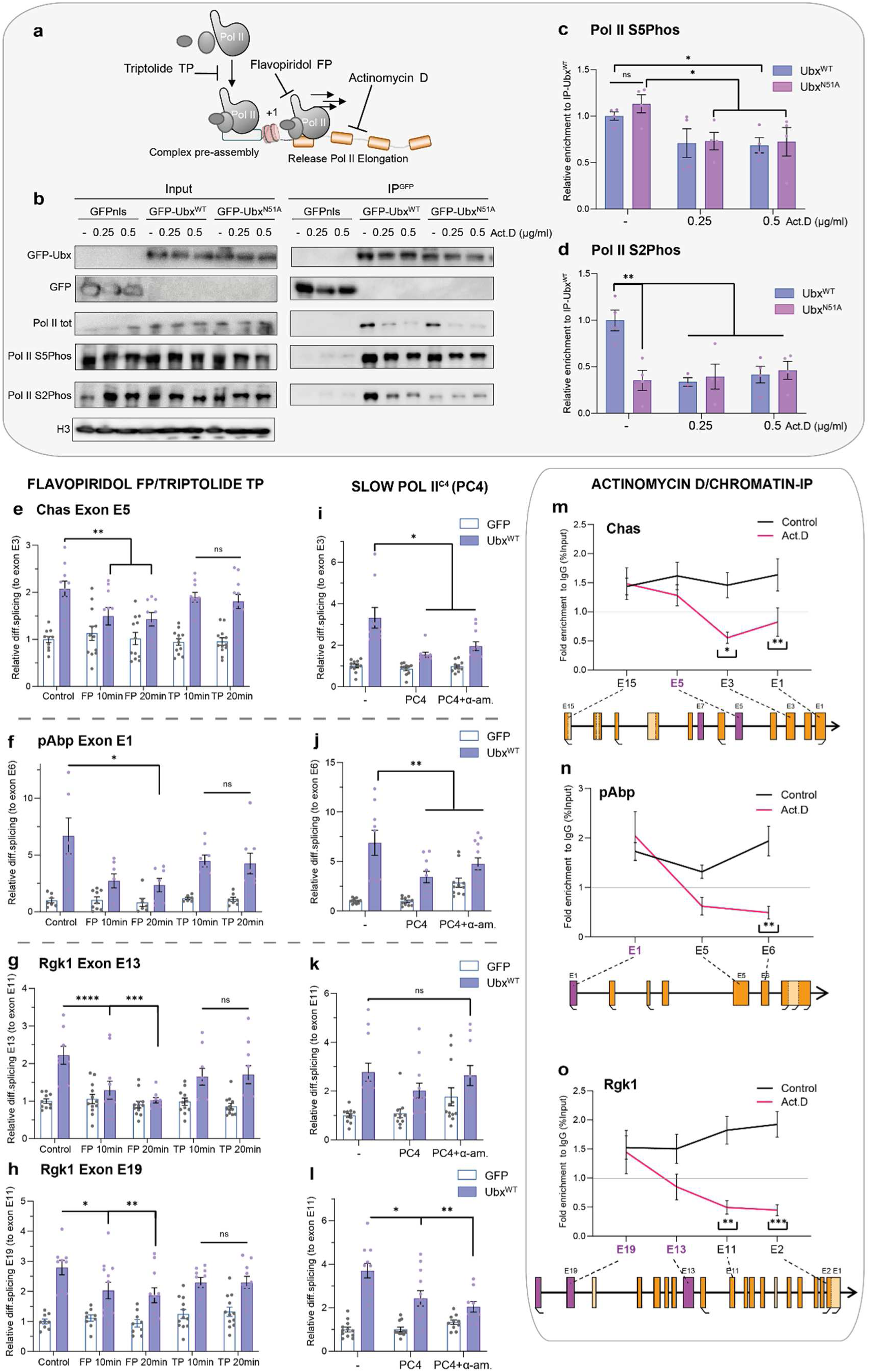
Ubx mediates co-transcriptional splicing via Pol II cooperation. **a.** Schematic of the inhibitory effect of transcription drugs, Flavopiridol (FP), Triptolide (FP) and Actinomycin D (act.D) on Pol II. **b.** Co-immunoprecipitation of Pol II total, paused (S5Phos) and active (elongating S2Phos) with GFP fusion proteins (GFPnls, GFP-Ubx^WT^, GFP-Ubx^N51A^), expressed in *Drosophila* S2R+ cells treated for 20h with actinomycin D (act.D) as indicated or DMSO as control (-). Western blots were probed with the indicated antibodies. The input is shown as a control of expression levels (lanes 1-9), Histone 3 (H3) is used as a loading control. **c-d.** Quantification of relative enrichment of Pol II S5phos (**c**) and S2Phos (**d**) relative to Ubx^WT^ and Ubx^N51A^ pull down showing once more a specific enrichment of active Pol II S2Phos for Ubx^WT^ compared to Ubx^N51A^, and a significant decrease of Pol II S5Phos (30%) and S2Phos enrichment (70%) upon actinomycin D treatment. n=4 independent biological replicates. Statistical test by one-way ANOVA (p<0.05*, p<0.01**, ns=non-significant). **e-l.** RT-qPCR experiments showing the differential retention of exon cassettes over constitutive exons for Chas (**e,i**), pAbp (**f, j**) and Rgk1 (**g-h, k-l**) in *Drosophila* S2R+ cells expressing GFP control, Ubx^WT^ (blue). **e-h**. 10 and 20 minutes treatments with Flavopiridol (FP, elongation repressed) or Triptolide (TP, elongation proceeds until termination) showed that Ubx activity on splicing depends on active transcription; (**i-l**) ectopic expression of the slow Pol II^C4^ (PC4) mutant characterised by a slow transcription rate and α-amanitin resistance, combined with α-amanitin treatment (α-am) showed that Ubx effect on splicing depends on the Pol II rate. **m-o.** Chromatin-ImmunoPrecipitation (ChIP) experiments of Ubx coupled with actinomycin D treatment are presented as percent of enrichment relative to input and IgG control (horizontal bar set to 1). Binding of Ubx on the proximal and distal exons to the Transcription Start Site (TSS) of Chas (**m**), pAbp (**n**) and Rgk1 (**o**) are displayed relative to a schematic of the genes architecture. Differentially spliced exons are highlighted in purple. n=3 independent biological in triplicates (RNA) or duplicates (ChIP). Bars represent mean±SEM. Statistical test by one-way ANOVA (p<0.05*, p<0.01**, p<0.001***, ns=non-significant). See also supplementary Fig.12.

To test how active transcription impacts on Ubx splicing activity, we analysed the effect of transcriptional drugs on Ubx splicing activity. The cells were treated with Flavopiridol (FP), an inhibitor of the Pol II kinase CDK9 which blocks the release of Pol II, thereby impairing elongation (Bensaude, 2011; McSwiggen et al., 2019). In parallel, cells were treated with Triptolide (TP), an inhibitor of TFIIH that prevents the assembly of the Pol II pre-initiation complex, while the engaged active Pol II can still achieve the ongoing transcription cycle (Bensaude, 2011; McSwiggen et al., 2019) (Fig.6a). Upon treatment with Flavopiridol, the differential splicing induced by Ubx^WT^ was significantly reduced for most of its target exon cassettes (Fig.6e-h, Supplementary Fig.12a,d). Flavopiridol specifically impaired the elongation process, indicating that Ubx splicing activity depends on active elongation. In contrast, the transcription inhibitor Triptolide had no effect on Ubx splicing activity on the selected target genes (Fig.6e-h, Supplementary Fig.12a,d). Of note, the differential splicing of the exon E4 of H3.3B was reduced, most likely due to its small gene size (Supplementary Fig.12a). This indicated that Ubx splicing activity depends on transcriptional elongation but not on initiation process.

We next reasoned that the rate of Pol II should impact on Ubx spliced target genes if Ubx regulates splicing co-transcriptionally. To test this hypothesis, we used a mutant of the biggest subunit of Pol II (rpII215), termed Pol II^C4^ (PC4) (Greenleaf, 1980; de la Mata et al., 2003). Pol II^C4^ is associated with a slower elongation rate than the Pol II^WT^ thereby impacting on splicing in a gene specific manner (Saldi et al., 2016). This mutant is resistant to α-amanitin, a drug inducing a degradation of the Pol II^WT^ form (Bensaude, 2011). Upon expression of Ubx in combination with Pol II^C4^ (PC4) and α-amanitin treatment in *Drosophila* S2R+ cells, we observed a gene-specific effect of the Pol II^C4^ on Ubx spliced targets (Fig.6i-l, Supplementary Fig.12d-e). Chas exon E5, pAbp exon E1 and Rgk1 exon E19 retentions were reduced while Rgk1 exon E13, Pura exon E11 splicing was not affected, and H3.3B exon E4 retention was significantly increased (Fig.6i-l, Supplementary Fig.12d-e). This indicated that Pol II elongation rate impacts on Ubx splicing activity in an exon-specific manner.

### Ubx dynamically binds chromatin during transcription

Our data indicated that the Ubx chromatin (or DNA) binding and Pol II interaction are necessary for its splicing activity. Moreover, Ubx regulates splicing co-transcriptionally and its interaction with active Pol II depends on its DNA-binding ability. Altogether, this strongly suggests that Ubx could regulate splicing via a dynamic interplay with the Pol II during active transcription, thereby binding or travelling along the gene body to regulate splicing. We sought to test this model by examining how active transcription affects Ubx binding dynamic within gene bodies. To this end, we performed Chromatin-ImmunoPrecipitation (ChIP) experiments of Ubx after actinomycin D treatment in *Drosophila* S2R+ cells (Fig.6m-o, Supplementary Fig.12f). We observed a significant enrichment of Ubx binding in exons proximal and distal to the Transcription Start Site (TSS-first exon) of Chas, pAbp, Rgk1 and Pura. Interestingly, Actinomycin D treatment significantly reduced the binding of Ubx in the distal exons of Chas (E3, E1), pAbp (E5, E6), Rgk1 (E13, E11, E2) and Pura (E5, E11, E14, E19), while proximal exons to the transcription start sites were still bound by Ubx similarly to control condition (Fig.6m-o, Supplementary Fig.12f). Notably, the binding of Ubx to the decapentaplegic *(dpp)* enhancer located in an intergenic region was not affected by the Actinomycin D treatment (Supplementary Fig.12g).

We next reasoned that Ubx dynamic should be affected by the Pol II speed rate if it travels along the gene body to regulate elongation-coupled splicing. To explore this possibility, we made use of Fluorescence Recovery After Photobleaching (FRAP) experiments to investigate the Ubx protein dynamics in cells (Fig.7a-c, Supplementary Fig.13). We compared the dynamic of GFP-Ubx^WT^ with GFP-Ubx^N51A^, the mutant which is no longer able to bind DNA, interact with active Pol II or coordinate splicing (Fig.7a-c). The dynamic behaviour of TFs has been largely linked to an exponential model with two-components (Phair et al., 2004). In detail, the model separates TFs population into i), fast mobile (diffusion and transient interaction), ii), slow mobile (scanning chromatin, longer interaction) and iii), immobile (stable interaction) fractions (Fig.7b). First, we confirmed the suitability of the mathematical model for Ubx dynamic by assessing the quality with the Akaike information criterion (AIC) and Bayesian information criterion (BIC). The lower value of the AIC and BIC for the double exponential compared to the single exponential models validated the suitability of the model for GFP-Ubx dynamic (Supplementary Fig.13a). Next, we assessed the distribution of GFP-Ubx populations revealing a larger immobile population of GFP-Ubx^WT^ compared to GFP-Ubx^N51A^ (Fig.7b). The immobile population thus refers to the Ubx^WT^ molecule stably bound to enhancers and promoters (Govindaraj et al., 2019). Notably, the residual immobile population observed for the GFP-Ubx^N51A^ could account for the protein clusters observed for both GFP-Ubx forms (Supplementary Fig.13c). Subsequently, we calculated the half-time recovery as well as the residence time of the GFP-Ubx proteins. Both values were smaller for Ubx^N51A^ than Ubx^WT^, indicating once more the loss of stable binding of Ubx^N51A^ to the chromatin (Fig7a, c, Supplementary Fig.13b).

**Figure 7:**
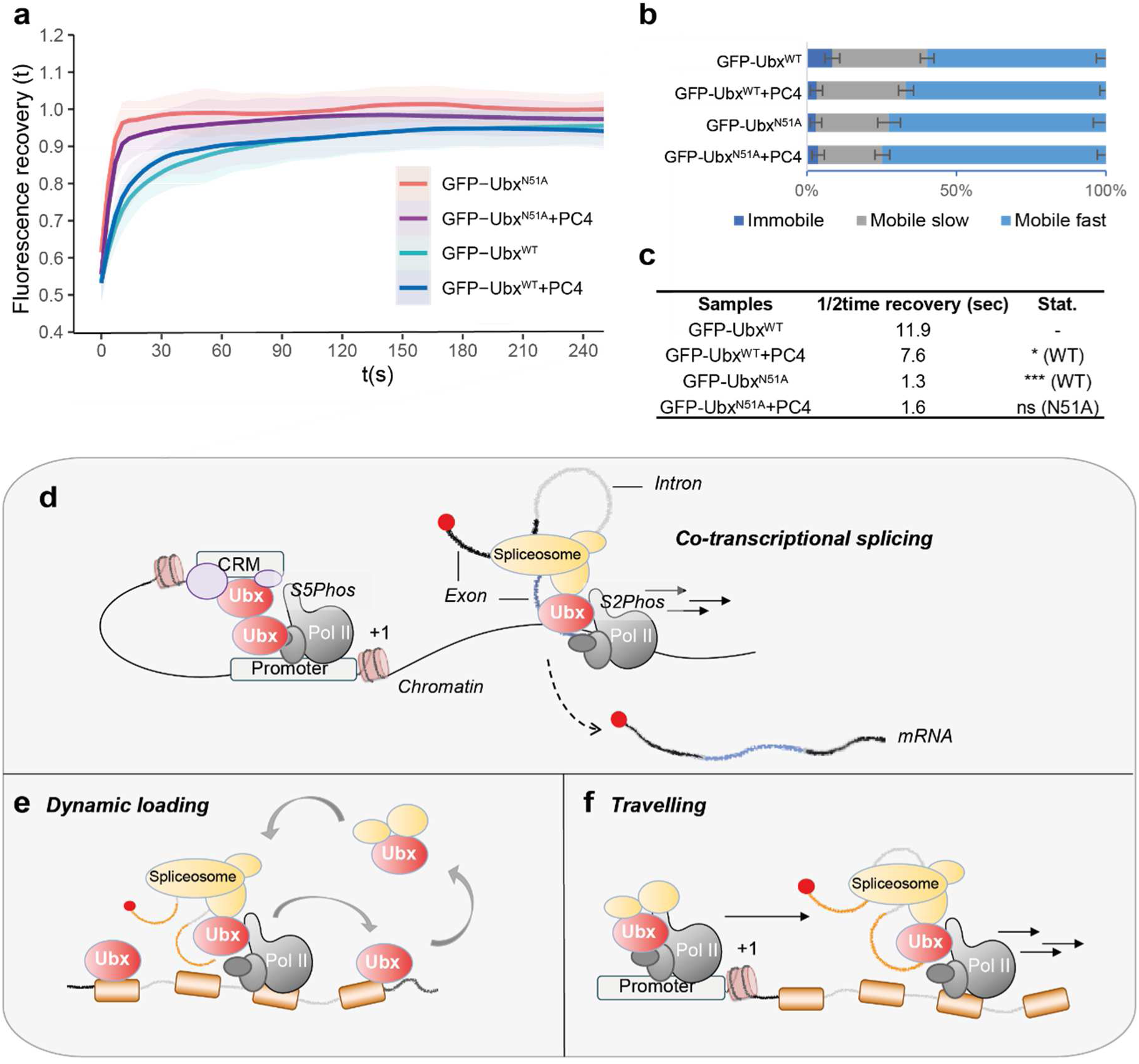
Models for Ubx regulation of co-transcriptional splicing and protein dynamics. **a.** Normalised curves of Fluorescence Recovery (t) After Photobleaching (FRAP) related to time (s, second) from data acquired using *Drosophila* S2R+ expressing GFP-Ubx^WT^ or GFP-Ubx^N51A^ co-expressed or not with slow Pol II^C4^ (PC4) coupled with α-amanitin treatment. First post-bleach acquisition data point is set to t(s)=0. Modelling and fitting are described in Materials and Methods. **b.** Distribution of Ubx populations is plotted for the different samples: immobile represents the fraction stably loaded onto the chromatin, slow mobile, the intermediate interactions, and fast mobile, the transient interaction as well as diffusion molecules of Ubx. Mean±SEM are shown. Statistical significance evaluated by t-test (p<0.05*). **c.** Value of the calculated half-time (1/2time) recovery and statistical differences compared to Ubx^WT^ condition are displayed. it shows that Ubx half time recovery is significantly faster for Ubx^N51A^ than Ubx^WT^. The slow Pol II^C4^ (PC4) reduces Ubx immobile population and Ubx^WT^ half time recovery, mirroring a potential decrease of stable binding of Ubx molecules on chromatin. n=17 for GFP-Ubx^N51A^, n=24 for GFP-Ubx^N51A^+PC4, n=21 for GFP-Ubx^WT^ and n=24 for GFP-Ubx^WT^+PC4 nuclei included in the analysis. **d-f.** Schematic of the proposed model for co-transcriptional splicing mediated by Ubx. **d.** Ubx regulates co-transcriptional splicing via a dynamic interplay with Pol II, the splicing machinery and a dual requirement of DNA-RNA interface. Two non-exclusive models for Ubx molecular mode of action are proposed: **e.** Ubx is dynamically loaded along the gene body relying on its DNA-binding abilities. Like a bouncing/scanning behaviour, Ubx recruits the spliceosome *de novo* on spliced exons which are recognized and bound via Ubx RNA-binding abilities. **f.** Ubx, paused Pol II and splicing factors are loaded together onto the promoter. Upon transcription activation, Ubx travels with the Pol II (S2Phos) and the splicing factors thereby scanning both the chromatin and RNA through different HD interface to allow the specific recognition and regulation of targeted exons. See also supplementary Fig.13.

In order to evaluate if the Pol II elongation activity affects Ubx dynamic, we co-expressed GFP-Ubx^WT^ in combination with the slow Pol II^C4^ (PC4) and α-amanitin. Surprisingly, the half-time recovery of GFP-Ubx^WT^ was significantly reduced in the presence of Pol II^C4^ (1/2t=11.9 to 7.6sec, Fig.7c). Moreover, the immobile fraction of Ubx^WT^ was reduced (8.5% to 3.2%) compared to control condition (Fig.7b). In contrast, the Ubx^N51A^ mutant was not affected by the Pol II rate (Fig.7a-c). This is in agreement with the interaction assay showing that Ubx^N51A^ interacts weakly with active Pol II (Fig.5a, S2Phos). Based on this result, we propose that i), the immobile fraction reflects Ubx stable binding on enhancer and promoter, ii), the slow mobile fraction refers to Ubx strong interaction or scanning the chromatin, iii), the fast mobile fraction is a combination of diffuse, and transiently bound molecules involved in the turn-over of transcription (i.e., initiation-termination). In this context, the slow polymerase reduces the half-time recovery of Ubx on the chromatin, as Ubx molecules are still bouncing or travelling along the gene body due to the slower transcription cycle (Fig.7d-f).

Altogether, the data indicated that Ubx dynamically binds to the chromatin during active transcription. This suggests that Ubx regulates co-transcriptional splicing via a dynamic interplay between chromatin, Pol II and RNA. From chromatin to mRNA and back, we anticipated various inter-connected mechanisms that will impact on Ubx co-transcriptional activities and ultimately, on the mRNA fate.

## DISCUSSION

Taken together, our studies uncovered a novel molecular function of the Hox TF Ubx in mRNA splicing. Ubx coordinates mRNA expression and splicing via common and distinct mode of actions, thereby promoting cell-type specific function in defined cell and tissue context. This extends the repertoire of Hox molecular function and its impact on cell fate. Our previous work revealed that Ubx interacts with a distinct set of regulatory cofactors in different embryonic tissues (Carnesecchi et al., 2020). The results further exhibited that Ubx interacts with several players of gene expression including mRNA splicing factors to coordinate tissue development (Carnesecchi et al., 2020). Altogether, this provides the molecular basis to explore the role of Ubx in mRNA splicing *in vivo* and a new perspective of Hox function for development and tissue homeostasis.

Our results indicated that the N51 amino-acid of the HD is essential for its splicing activity but still mediates partly transcriptional regulatory function by repressing and activating genes expression. First, this could be due to residual chromatin binding via protein interactions. ChIP-Seq experiments performed in *Drosophila* Kc167 cells with a combinatorial mutated version of Ubx HD showed that Ubx mutant retained few binding to the chromatin (Beh et al., 2016). Nonetheless, the FRAP experiments indicated that the punctual mutation N51A is essential for the stable binding of Ubx to the chromatin (Fig.7). Second, the effect could be due to a loss of interaction with splicing factors. However, our previous study revealed that Ubx^N51A^ protein is still able to interact with splicing factors in cells and in embryos (Carnesecchi et al., 2020). Importantly, the N51A mutation impairs Ubx DNA-binding ability, but is not essential for RNA-binding of Ubx *in vitro*. Thus, the effect mediated by Ubx^N51A^ on Ubx target genes expression could be due to its ability to bind mature mRNA in the cytoplasm. In this context, Ubx could have additional RNA-binding specific function, such as regulating RNA transport, decay or translation as suggested for other Hox proteins (Rezsohazy, 2014; Topisirovic et al., 2005). We showed previously that ectopic expression of Ubx^N51A^ is not able to induce homeotic transformation in embryos (Carnesecchi et al., 2020). However, Ubx^N51A^ partly shapes the transcriptional (and not the splicing) program in the Hox-free S2R+ *Drosophila* cells. Analysing the impact of Ubx^N51A^ on homeosis and morphogenesis may provide key information about the transcriptional and post-transcriptional functions of Ubx in development (Hombría and Lovegrove, 2003).

Our study shed light on a novel ability of Ubx to bind RNA *in vivo* and *in vitro*. This is defining Ubx as member of the DNA- and RNA-binding proteins (DRBPs), together with the Sox TFs or the HD-TF Bicoid (Bcd) (Cassiday, 2002; Hudson and Ortlund, 2014; Niessing et al., 2000). Ubx binds several RNA exonic sequences *in vivo* and *in vitro*. In contrast, the RNA-binding ability of Bcd is restricted to *Caudal,* for which Bcd binds a putative RNA sequence in the 3’UTR (Rivera-Pomar et al., 1996). This depends on the amino acid R54 of Bcd HD, which is not conserved for Ubx (Supplementary Fig.14). Therefore, Ubx most likely uses a different HD- RNA interface.

The largest family of TFs described so far for having dual abilities of DRBPs and a role in mRNA splicing is the Sox group (Girardot et al., 2018; Holmes et al., 2020; Hou and Yu, 2010; Ohe et al., 2002; Rambout et al., 2018). The Sox and HD family are both member of the Helix-turn-helix (HTH) DNA-binding domain containing TFs (Auer et al., 2020), sharing similarity of sequences (Supplementary Fig.14). Interestingly, Sox2 binds the long coding RNA ES2 in a non-sequence specific manner (Holmes et al., 2020). In contrast, Ubx seems to employ a specific RNA-recognition interface, as Ubx binding *in vitro* was not detected for the pAbp exon E6. Yet, the results cannot account for binding events happening on non-labelled UTP-free sequences. In contrast, the difference was clearly mitigated for Chas exon E3 and E5. This provides striking clues of a Hox-RNA paradigm: Ubx binds RNA *in vitro* with less specificity than *in vivo.* Thus Ubx-RNA specificity could be mediated *in vivo* by the interaction with cell-type specific splicing factors as previously emphasised (Carnesecchi et al., 2020). Moreover, it is most conceivable that the RNA structure and shape will be an essential parameter for investigating the RNA-binding affinity of Ubx in future.

We found that the amino acid N51 of Ubx is non-essential for its *in vitro* RNA-binding properties. Interestingly, none of the amino acids of Sox2 involved in DNA-specific contact are essential for its RNA-binding (Holmes et al., 2020). The DNA-binding domain of Sox2 is sufficient to mediate the interaction, while engaging DNA and RNA with a different interface (Holmes et al., 2020). In this context, how does Ubx mediate RNA-DNA interaction? Is it mutually exclusive? Is Ubx dimerized for contacting both RNA and DNA molecules? Furthermore, our results indicated that the Ubx-HD mediates a different binding affinity compared to the full-length protein. Sox2 also mediates RNA-binding via an additional 60 amino acid RNA-binding motif (RBM) (Hou et al., 2020). Even more important, the RBM domain of Sox2 provides RNA sequence specificity as shown by SELEX experiments. Altogether, it suggests that Ubx-RNA binding affinity relies on multiple interaction interfaces.

In this study, we provide the first evidence for a dynamic interplay between Ubx and Pol II to regulate splicing. First, our data indicate that Ubx interacts with Pol II, both with the paused (S5Phos) and active (S2Phos) forms. This is consistent with a large amount of studies showing that Ubx and the Hox TFs in general regulate transcription activation (Johnson and Krasnow, 1990; Mortin and Lefevre, 1981; Mortin et al., 1992) promoter pausing, Pol II release (Choe et al., 2009; Chopra et al., 2009; Zouaz et al., 2017), and interact with various components of the pre-initiation complex (Boube et al., 2014; Prince et al., 2008). Yet, no direct interaction between Hox and Pol II were identified. Our results show a direct interaction between Ubx and the conserved CTD of Pol II. How this interaction is mediated *in vivo* and impacts on Ubx transcriptional and splicing activity is still to explore.

Second, using transcriptional drugs and mutant of the Pol II, our results revealed that Ubx mediates elongation-coupled splicing. Interestingly, the rate of the Pol II affects Ubx spliced target genes differently. Thus, Ubx may regulate splicing via various molecular mechanisms, such as by promoting the recruitment of splicing activators (SR proteins) or repressors (hnRNP), by modifying the RNA folding or impacting on the chromatin landscape (Carrocci and Neugebauer, 2019; Saldi et al., 2016, Carnesecchi et al., 2018). Taken together, this further extend the possible combination of Hox molecular mode of actions.

Third, ChIP and FRAP experiments revealed that Ubx chromatin binding and protein dynamic are affected by the transcription rate. Interestingly, the recovery rate of Ubx was not extended in presence of slow Pol II^C4^. This suggests once more that Ubx uses different mechanisms to regulate splicing, within one of them in close-cooperation with the Pol II.

Altogether, the data suggests a model in which Ubx regulates co-transcriptional splicing via a dynamic interplay with Pol II, the splicing machinery and a dual requirement of DNA-RNA interface. We proposed herein two non-exclusive models for Ubx molecular mode of action (Fig.7d-f). First, Ubx is dynamically loaded along the gene body, with a binding/bouncing behaviour, thereby allowing the recruitment of the spliceosome machinery for its *de novo* assembly (Fica and Nagai, 2017). Second, Ubx is initially loaded onto the chromatin in complex with Pol II and splicing factors at the promoter. Upon transcriptional activation, they travel together to regulate transcription and splicing. In future, single molecule imaging strategy will be crucial to investigate these co-transcriptional regulatory models (Lerner et al., 2020).

All in all, our work lays the groundwork to understand the role of Hox proteins in mRNA splicing, thereby providing new perspectives of Hox function in development and diseases. Beyond the Hox TFs, it broadens our insights into the molecular mechanisms employed by TFs to coordinate the variety of cell and tissue identities.

## MATERIALS AND METHODS

### Fly line and vector constructs

The following fly lines were used for the study: *GFP-Ubx* (Domsch et al., 2019), *w1118.* The PC4 plasmid was kindly obtained from the Drosophila Genomic Research Center (DGRC). The Ubx plasmids were generated previously for (Carnesecchi et al., 2020). The vector constructs used in the study are pActin-Gal4, PC4 (from DGRC), UAS-GFPnls, UAS-myc-Ubx^WT^, UAS-myc-Ubx^N51A^, UAS-GFP-Ubx^WT^, UAS-GFP-Ubx^N51A^, pET-his-Ubx^WT^, pET-his-Ubx^N51A^, pET-MBP-his-HD^WT^, pET-MBP-his-HD^N51A^, pGEX-Ubx-FL, pGEX-Ubx-Nter, pGEX-Ubx-Cter, pGEX-Ubx-HD, pGEX-GST, pActin-BetaGalactosidase for plasmid quantity normalisation. For protein-RNA assay, probes listed in Supplementary Table 7 were cloned in pGEM®-T Easy (Promega) for T7 polymerase transcription.

### Cell culture and transfection

*Drosophila* S2R+ cells were maintained at 25°C in Schneider medium supplemented with 10% FCS, 10U/ml penicillin and 10µg/ml streptomycin. Cells were simultaneously seeded and transfected with Effectene (Qiagen) according to the manufacturer’s protocol. For all constructs the Gal4-UAS system was used for inducible protein expression driven by the Actin promoter. For whole cell protein extracts, cells were harvested in Phosphate Buffered Saline (PBS) and pellets were resuspended in RIPA buffer supplemented with protease inhibitor cocktail (Sigma-Aldrich). For RNA analysis, cells were seeded in 6 well plates and transfected as described with UAS-GFPnls, UAS-myc-Ubx^WT^, or UAS-myc-Ubx^N51A^ and pActin-Gal4. For interaction, ChIP and RIP assay, 10.10^6^ cells were seeded in 100mm dishes and transfected as described with UAS-GFPnls, UAS-GFP-Ubx^WT^, or UAS-GFP-Ubx^N51A^ and pActin-Gal4. Cells were harvested in Phosphate Buffered Saline (PBS) after 48 hours of transfection and pellets were resuspended with lysis buffer supplemented with protease inhibitor cocktail (Sigma-Aldrich) and 1 mM of DTT. For ChIP and interaction experiments, 0.25 or 0.5 µg/ml of Actinomycin D treatment (Sigma-Aldrich) was applied for 20 hours. For Pol II^C4^ (PC4) expression experiments, 5 µg/ml α-amanitin (Sigma-Aldrich) treatment was applied for 25-30 hours before RNA extraction or FRAP analysis. 10 µM Triptolide (Sigma-Aldrich) and 10 µM Flavopiridol (Sigma-Aldrich) treatment were applied for 10 and 20 minutes. For FRAP analysis, cells were seeded and transfected in 12 well plates and transferred with fresh supplemented media (supplemented with α-amanitin for PC4 experiment) in glass bottom dishes coated with Poly-lysine (Sigma), at least 2 hours before image acquisition.

### RNA-Seq from *Drosophila* S2R+ cells

Total RNAs were extracted from four independent replicates from *Drosophila* S2R+ cells expressing GFPnls, myc-Ubx^WT^, or myc-Ubx^N51A^ (Gal4-UAS, actin promoter) using Qiagen RNA extraction kit (RNeasy). RNA quality was assessed using BioAnalyzer 2100TM (Agilent Technologies). Material handling and mRNA-Seq directional libraries were performed with the Deep-Sequencing facility in Heidelberg (Cell Networks). Sequencing was performed with NextSeq500 High-Output with a read-length of 75bp and single-end strands. Replicates were validated by FastQC report and 3 replicates for each sample were further selected according to Principle Component Analysis (PCA) analysis for the study.

### Expression and differential splicing analysis of RNA-Seq

RNA-Seq analysis was performed by using genome 6 (dm6). The quality of the RNA-Seq reads was quantified via FastQC. Trimming was performed with java script “Trimmomatic 0.36”(Bolger et al., 2014), for the removal of the adapter (TruSeq-SE sequencing adapter), with the following command line: java -jar /path_to_java_script/Trimmomatic- 0.36/trimmomatic-0.36.jar SE -phred33 <file.txt> <outputfile.txt> ILLUMINACLIP:TruSeq- SE.fa:2:30:10 LEADING:3 TRAILING:3 SLIDINGWINDOW:4:15 MINLEN:36. The reads were aligned using STAR (genome generated: STAR --runThreadN 14 --runMode genomeGenerate --genomeFastaFiles Drosophila_melanogaster.BDGP6.dna.toplevel.fa --sjdbGTFfile Drosophila_melanogaster.BDGP6.86.gtf --genomeSAindexNbases 12; running STAR: STAR --runThreadN 14 --genomeDir /STAR/GenomeDir/ --readFilesIn /STAR/ <file> --outSAMtype BAM Unsorted) (Dobin et al., 2013). Aligned reads were counted by HTSeq (htseq-count -f bam -r pos -m union -s no -t exon accepted_hits.bam Drosophila_melanogaster.BDGP6.86.gtf > count.txt) for further analysis in DESeq (Anders et al., 2015). For the analysis in Junction-Seq the aligned reads were counted and the genome flattened with QoRTs by using the available java scripts (counting: java -jar /path/QoRTs.jar QC -- singleEnded --minMAPQ 75 --nameSorted sorted.bam Drosophila_melanogaster.BDGP6.86.gtf rawCts/file; flattened: java -jar /path/QoRTs.jar makeFlatGff --stranded Drosophila_melanogaster.BDGP6.86.gtf annoFiles/JunctionSeq.flat.gff.gz) (Hartley and Mullikin, 2015). Differential expression analysis was performed with DESeq2 under standard conditions (Love et al., 2014). Different exon usage and the identification of splicing variants was identified by using the R tool Junction-Seq under standard conditions (Hartley and Mullikin, 2016). All data results were further processed in Excel and False Discovery Rate (FDR) was established at FDR 0.1.

### ChIP-Seq genome wide distribution

Analysis of Ubx DNA-binding profile in *Drosophila* S2 cells (Zouaz et al., 2017) and in the mesoderm (Domsch et al., 2019) where performed as in Domsch et al. (2019). The aligned read files (BAM) of the Ubx ChIP-Seq in *Drosophila* S2 cells (Zouaz et al., 2017) and in the differentiating mesoderm (Domsch et al., 2019) were downloaded from GEO (accession GSE101556 and GSE121754 respectively). Peak calling was performed using MACS2 with standard narrow peak settings (macs2 callpeak -t ubx_aligned_reads.bam -c input_aligned_reads.bam -g 1.28e8 -f BAM --outdir output_dir -n ubx -q 0.01). Statistics and annotation of the called peaks was performed using HOMER annotatePeaks.pl (annotatePeaks.pl ubx_peaks.narrowPeak dm6 -annStats ubx_stats.txt > annotated_mlUbx_chr_peaks.txt).

### RNA extraction and quantitative PCR

Total RNAs for 3 to 4 independent experiments performed in triplicate were extracted by Trizol (Ambion) according to the manufacturer’s protocol. We converted 1μg of RNA to first strand cDNA using Reversaid kit (ThermoFisher Scientific) and random hexamers. Real-time PCR was performed in a 96-well plate using the Platinum™ SYBR™ Green (Invitrogen). Data were quantified by the ΔΔ-Ct method and normalised to Actin 5C expression or internal region of constitutively expressed exons of the related gene. Sequences of the primers used in this study are provided in Supplementary Table 8 and efficiency was confirmed with cDNA serial dilution.

### RNA-Immunoprecipitation and quantitative PCR

Confluent *Drosophila* S2R+ cells expressing GFPnls, GFP-Ubx^WT^ or GFP-Ubx^N51A^ plated in 100mm dishes were collected in cold PBS 48h after transfection. After several PBS washes, cell pellets were resuspended in buffer A (10 mM Hepes pH 7.9, 10 mM KCl, 1.5 mM MgCl2, 0.34 M sucrose, 10% glycerol). Lysates were incubated with 0.1% Triton and centrifugated. Nuclear pellets were then resuspended with IP Buffer (150mM NaCl, 50mM Tris pH 7.5, 1mM EDTA, 1% Triton), incubated on ice with vigorous regular vortex and sonicated (8x 30 sec on/off, Picoruptor, Diagenode). All buffers were supplemented with protease inhibitor cocktail (Sigma), 1 mM of DTT, 0.1 mM PMSF and RNAsine (Promega). Input fractions were collected, both for protein and RT-qPCR control. 1.5-2 mg of nuclear lysates were diluted in IP buffer (20 mM Tris pH7.5, 150 mM NaCl, 2 mM EDTA, 1% NP40) and incubated for 4 hours with 40 µl of GFP-Trap beads (Chromotech). Beads were washed 5 times for 5 min with IP buffer. 10% were collected for protein analysis, the remaining beads were resuspended in Trizol and RNAs were extracted according to the manufacturer’s protocol. Notably, RNAs were precipitated for 1 hour at -80°C in isopropanol to increase efficiency. Retro-transcriptions were proceed as described, on 2 µg of RNA by doubling the total volume. cDNAs were then diluted by two and 2µl were used for quantitative qPCR. Enrichment was calculated relative to input and GFP values.

### Chromatin-Immunoprecipitation coupled with quantitative PCR

48h post-transfection, confluent *Drosophila* S2R+ cells expressing GFPnls, GFP-Ubx^WT^ or GFP-Ubx^N51A^ plated in 100mm dishes were crosslinked with 1% formaldehyde and quenched them for 5 min in 0.125 M Glycine. After several PBS washes, cell pellets were resuspended in lysis buffer (1% SDS, 50 mMTris·HCl, pH 8, 10 mM EDTA). Sonication (10 cycles, 30sec on/off) was performed with Picoruptor (Diagenode) according to the Diagenode recommendation. Lysates were incubated with 5 μl of antibody against Ubx (Guinea-pig, home-made) or IgG (santa cruz) overnight at 4 °C on rotation and for one additional hour with mixed Dynabead protein G and A (20:10 µl, Life Technologies). Beads were washed with TSE- 150 (0.1% SDS,1% Triton, 2 mM EDTA, 20 mM Tris, pH 8.1, 150 mM NaCl), TSE-500 (as TSE-150 with 500 mM NaCl), LiCl detergent (0.25 M LiCl, 1% Nonidet P-40, 1% Sodium Deoxycholate, 1 mM EDTA, 10 mM Tris, pH 8.1), and Tris-EDTA (5:1 mM). Combined elution and decrosslinking were performed by adding RNAse for 30min at 37°C, then 0.1% SDS with proteinase K for 1 hour at 37°C followed by additional incubation with NaCl under 900rpm shaking for 7 hours at 65 °C. DNA fragments were purified using Qiaquick minielute (Qiagen) according to the manufacturer protocol and diluted to 1/10 for input and to the half for immunoprecipitated fractions. qPCRs were performed using 2μL of DNA, and enrichment was calculated relative to input and IgG values.

### Co-immunoprecipitation of whole cell lysate and embryos nuclear extract

For co-immunoprecipitation assays, *Drosophila* S2R+ cells expressing GFPnls, GFP-Ubx^WT^ or GFP-Ubx^N51A^ were harvested in Phosphate Buffered Saline (PBS) and pellets were resuspended in NP40 buffer (20mM Tris pH7.5, 150mM NaCl, 2mM EDTA, 1% NP40) and treated with Benzonase (Sigma). GFP-Trap beads (Chromotek) were added to the protein 1.5- 2 mg of protein extract, incubated for 3 hours and washed 5 times with NP40 buffer.

For *in vivo* interaction from embryos, overnight collection of embryos was dechorionated, fixed with 3.2% formaldehyde and collected in PBS supplemented with Tween 0.1%. Pellets were resuspended in buffer A (10 mM Hepes pH 7.9, 10 mM KCl, 1.5 mM MgCl2, 0.34 M sucrose, 10% glycerol) and dounced 25-30 times with loose- and 5 times with tight-pestle. Lysates were filtered, incubated with 0.1% Triton and centrifugated. Nuclear pellets were then resuspended with buffer B (3mM EDTA pH8, 0.2mM EGTA pH8), sonicated (Picoruptor, Diagenode), and treated with Benzonase. 4 to 5 mg of nuclear lysates were diluted in NP40 buffer (20 mM Tris pH7.5, 150 mM NaCl, 2 mM EDTA, 1% NP40) and incubated overnight with 40 µl of GFP-Trap beads (Chromotek). Beads were then washed 5 times with NP40 buffer and all samples were resuspended in Laemmli buffer for immunoblotting analysis. All buffers were supplemented with protease inhibitor cocktail (Sigma), 1mM of DTT and 0.1mM PMSF. Input fractions represent 1-10% of the immunoprecipitated fraction.

### SDS-Page and Immunoblotting

For western blot analysis, proteins were resolved on 8 to 15% SDS-PAGE, blotted onto PVDF membrane (Biorad) and probed with specific antibodies after saturation. The antibodies (and their dilution) used in this study were: Ubx (home-made, 1/200), Histone 3 (Abcam, 1/10,000), GFP (Life Technologies, 1/3000), Tubulin (Serotec, 1/2000e), GST (Cell signalling, 1/5000e), Pol II total (Bethyl, A300-653A, 1/2000e), Pol II S5Phos (Bethyl, A304-408A, 1/1000e), Pol II S2Phos (Bethyl, A300-654A, 1/1000e).

### Protein purification and GST-Pull down

His-tagged and GST-tagged proteins were cloned for this study or from our previous work (Carnesecchi et al., 2020) in pET or pGEX-6P plasmids respectively. His- and GST-tagged proteins were produced from BL-21 (RIPL) bacterial strain, purified on Ni-NTA agarose beads (Qiagen) or Gluthatione-Sepharose beads (GE-healthcare) respectively and quantified by Coomassie staining. His-tagged proteins were specifically eluted from the beads with Imidazole. *In vitro* interaction assays were performed with equal amounts of GST or GST fusion proteins in affinity buffer (20mM HEPES, 10μM ZnCl_2_, 0.1% Triton, 2mM EDTA) supplemented with NaCl, 1mM of DTT, 0.1mM PMSF and protease inhibitor cocktail (Sigma). Proteins produced *in vitro* were subjected to interaction assays for 2 hours at 4°C under mild rotation with *Drosophila* nuclear extracts or recombinant human CTD (Active Motif). Bound proteins were washed 4 times and resuspended in Laemmli buffer for western-blot analysis. Input fraction was loaded as indicated.

### *In vitro* transcription with Cy3-UTP labelling

For *in vitro* transcription, selected fragments of alternatively spliced exons (Supplementary Table 7) were cloned into the pGEM®-T Easy Vector (Promega). To generate the DNA templates for transcription, plasmids were amplified in DH5α bacterial strain, purified and linearized 3’ to the cloned sequence using the SpeI restriction site. For producing internally labelled RNA, *in vitro* transcription was performed using the HighYield T7 Cy3 RNA Labelling Kit (Jena Bioscience, RNT-101-CY3) in accordance to the instructions of the manufacturer. Each reaction contained 500 ng DNA template, 0.4 μl UTP-X-Cy3 (5 mM) and 0.4 μl RiboLock RNase Inhibitor (ThermoFisher) and was incubated for 1 hour at 37°C. DNA template was digested with 1 μl TURBO™ DNase (ThermoFischer) for 15 min at 37°C. Finally, labelled RNA probes were purified using the ProbeQuant™ G-50 Micro Columns (Sigma) and eluted in 50 μl.

### Protein-RNA UV-crosslinking assay

To prepare the RNA–protein complexes for UV-crosslinking, 2 pmol of internally labeled RNA probes were mixed with approximately 0.5-1 μg of His-purified proteins. The binding reaction was performed in a pre-cooled 96-wells-plate in a volume of 30 μl containing 1x Binding Buffer (20mM Hepes pH 7.9, 1.4mM MgCl2, 1mM ZnSO4, 40mM KCl, 0.1mM EDTA, 5% Glycerol), 2 μg tRNA (ThermoFischer), 3 μg BSA, 10mM DTT and 0.1% NP40. After 20 min on ice, the samples were irradiated with UV light in a Stratalinker® UV Crosslinker Model 1800 (Stratagene) for 10 min on ice and subsequently transferred in Eppendorf Tubes. 1.5 μl of RNase A (10 mg/ml) (ThermoFischer) were added and the samples were incubated for 20 min at 37°C. Cy3-labelled protein-RNA complexes were resolved on 12% SDS-PAGE for 1 hour at 200 V and detected by fluorescence using INTAS Imager. Following the detection, the gels were stained with Coomassie overnight and imaged using a conventional image scanner (Epson).

### DNA-EMSA

The 5’-Cy5 labelled complementary oligonucleotides (Eurofin) commercially produced were annealed before reaction. The sequences used for this study were the following: Ubx sites: Cy5-5’-TTCAGAGCGAATGATTTATGACCGGTCAAG-3’. The binding reaction was performed for 20 min in a volume of 30μl containing 1x Binding Buffer (20mM Hepes pH 7.9, 1.4mM MgCl2, 1mM ZnSO4, 40mM KCl, 0.1mM EDTA, 5% Glycerol), 0.2μg Poly(dI-dC), 0.1μg BSA, 10mM DTT and 0.1% NP-40. For each reaction His-purified proteins were used. Separation was carried out (200V, 50min) at 4°C on a 4-6% acrylamide gel in 0.5x Tris-borate-EDTA buffer to visualize complex formation by retardation. Cy5-labelled DNA-protein complexes were detected by fluorescence using INTAS Imager.

### Fluorescence Recovery After Photobleaching acquisition and modelling

Timeseries were acquired on a NIKON A1R CSLM equipped with a 63x, NA 1.27, WI objective. Detection was done on a Galvano scanner at 64x64 px. The first 10 pre-bleach frames and 10 post-bleach frames were recorded at 33 frame per second (fps). Subsequent recovery was measured at 30 fps, followed by 0.2 fps for 80 frames, totalling about 4.5 minutes. Half nuclei were bleached for 250 ms at 100% laser power.

Acquired time series were analysed in ImageJ. Stacks were aligned using the Template Matching and Slice Alignment plugin. Total nuclear intensity (Inuc), half-nucleus bleached intensity (Ibl), and background intensity (Ibg) were used for analysis in R.

FRAP recovery was double normalised as follows: FRAP(t)=(I_nuc-I_bg)/(I_bl-I_bg)

FRAP_norm (t)=(FRAP(t))/I_pre

FRAP_(double“_” norm)=(FRAP_norm (t)-FRAP_norm (t_bl))/(1-FRAP_norm (t_bl))

The double normalized recovery curves were then fitted with the minpack.lm package to single and double exponential diffusion models:

Single: F(t)=M_mob-M_mob⋅e^(-k⋅t) t-half : t_(1/2)=-(lnα(0.5)/k) Immobile fraction: M_imb=1-M_mob

Double: F(t)=(M_fast+M_slow)-M_fast⋅e^(-k_on⋅t)-M_slow⋅e^(-k_off⋅t)

t-half: calculated using the investr package

Immobile fraction: M_imb=1-M_fast-M_slow

### Data analysis and visualisation

For GO-Term annotations and overrepresented GO-Term related to biological process analysis were performed with the web-tools PANTHER using Fisher test and FDR correction.

Integrative Genomics Viewer (IGV version 2.8.12) was used for Ubx ChIP-Seq data visualisation at logarithm scale using Bedgraph files.

Gel quantifications were performed with Fiji (is just image j).

Statistical analyses were performed using one-way Anova, Chi^2^ for distribution and t-test multiparametric using GraphPad Prism 9 software.

Data visualisation was achieved with GraphPad Prism 9, Microsoft office power point, excel and Adobe Illustrator.

### Data deposition

Raw data of RNA-Seq performed in *Drosophila* S2R+ cells will be deposited on Gene Expression Omnibus.

### Data accessibility

Embryonic mesoderm Ubx ChIP-Seq (GSE121754, mesoderm late) and RNA-Seq (GSE121670) were from (Domsch et al., 2019). Ubx ChIP-Seq (GSE101556) in *Drosophila* S2 cells are from (Zouaz et al., 2017).

## ACKNOWLEDGEMENTS

We thank the Bloomington Center for fly lines, and DGRC for plasmids. We would like to further thank Alexandra Moreira for sharing PTB plasmid, Jean-Yves Roignant for the RIP protocol, Guido Grossmann and Jan Lohmann for access to critical equipment. We thank the Nikon Imaging Center for access to FRAP facilities as well as the deep sequencing facilities of the Cell Networks. We are very grateful for the people who helped to improve the manuscript in particular Samir Merabet. JC is supported by the Fondation pour la Recherche Médicale (ARF202004011788). This project was supported in part by DFG LO 844/8-1 (IL).

## CONTRIBUTION

Conceptualisation: JC, PP. Experimental Design: JC, PP. Experimental procedures: JC, PB. FRAP acquisition and modelling: PNS. Bioinformatic data curation: KD, HP. Data analysis: JC. Supervision: JC. Writing, editing: JC with significant support of all authors. Funding acquisition: IL.

## ETHIC DECLARATIONS

The authors declare no competing interests

## Supplementary figures

See supplementary file

