## Supplementary figures for "The Hox transcription factor Ultrabithorax binds RNA and regulates co-transcriptional splicing through an interplay with RNA polymerase II"

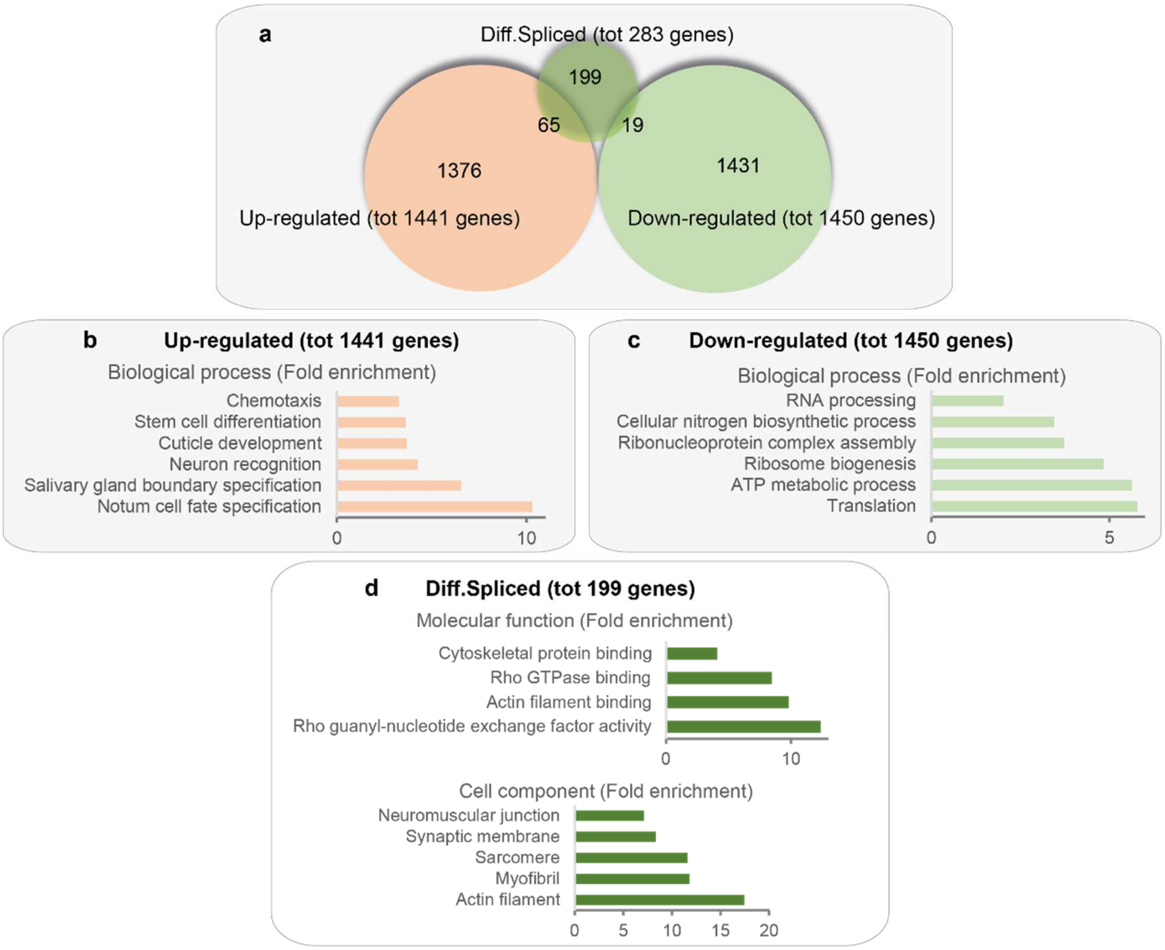


**Supplementary Figure 1: Transcriptome from *Drosophila* embryonic mesoderm upon degradation of Ubx. a.** Venn diagram overlapping the up-regulated (i.e., repressed, 1441), down-regulated (i.e., activated, 1450) and differentially spliced (283) genes upon Ubx depletion in the embryonic mesoderm (Domsch et al., 2019). **b-c.** Fold enrichment of Gene ontology (GO) term of biological processes enriched for the list of genes up-regulated (1441) and down-regulated (1450) upon Ubx mesoderm degradation (pvalue<0.05, fold enrichment). **d.** Fold enrichment of Gene ontology (GO) term of molecular function (upper panel) and cell component (lower panel) enriched for the list of genes only differentially spliced (199/283) upon Ubx depletion (pvalue<0.05, fold enrichment).

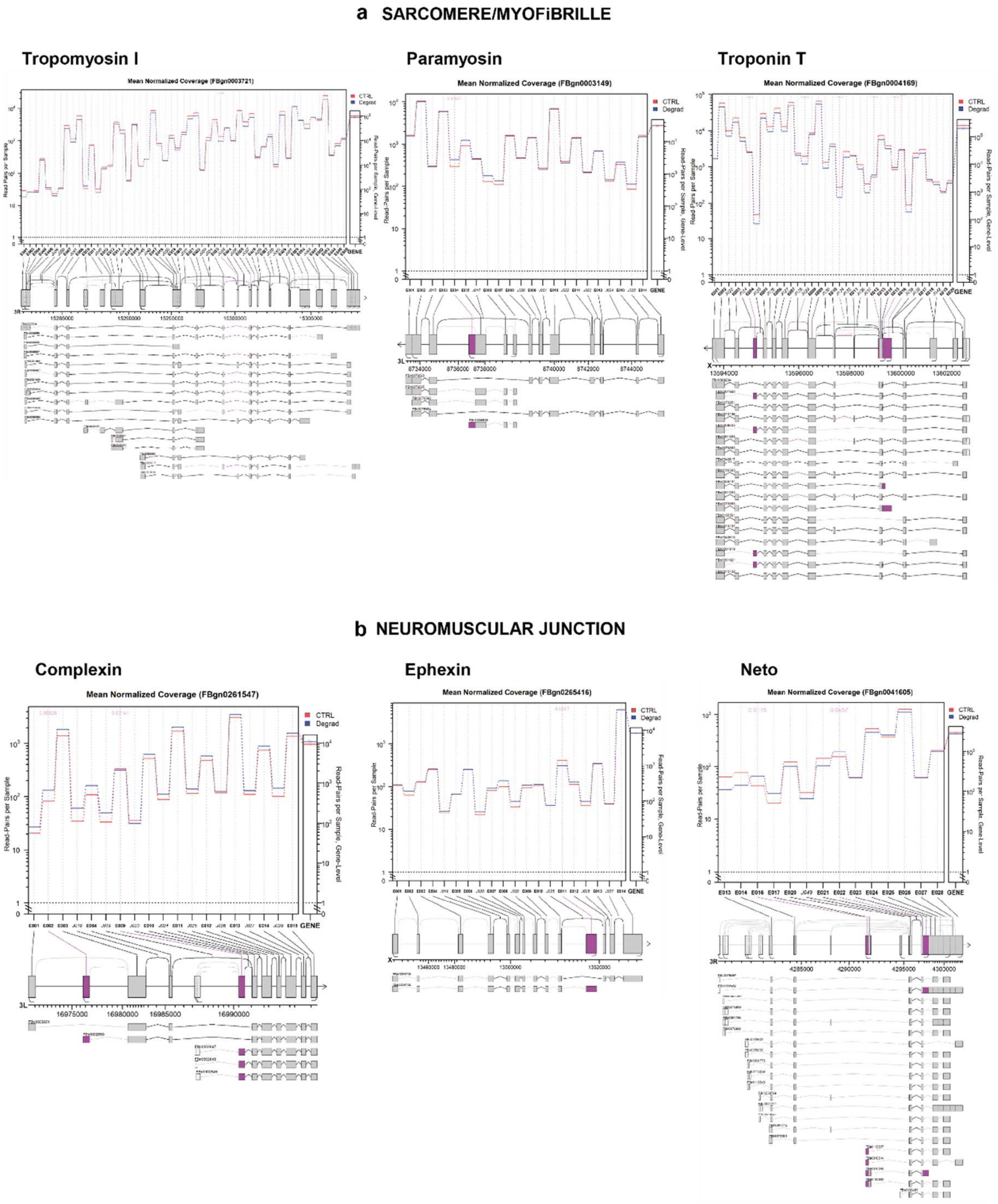


**Supplementary Figure 2: Ubx degradation impacts on mesoderm-specific alternatively spliced genes in *Drosophila* embryos. a-b.** Representative genes involved in (**a**) myofibrille/sarcomere modules (Tropomyosin I, Paramyosin, Troponin T) and (**b**) neuromuscular junction (Complexin, Ephexin, Neto), differentially spliced in embryonic mesoderm upon Ubx depletion (Domsch et al., 2019). Visualisation from Junction-Seq of the mean normalised read count for each exon or splice junction (left Y-axis, extended panel), and the gene level normalised read count (right Y-axis, narrow panel) of genes differentially spliced upon Ubx depletion in the mesoderm (degrad, blue line) compared to control (CTRL, red line).

Significant differential splicing events are highlighted in purple. Isoforms including differentially spliced exon or junction usages (purple) upon Ubx degradation are displayed below each read count.

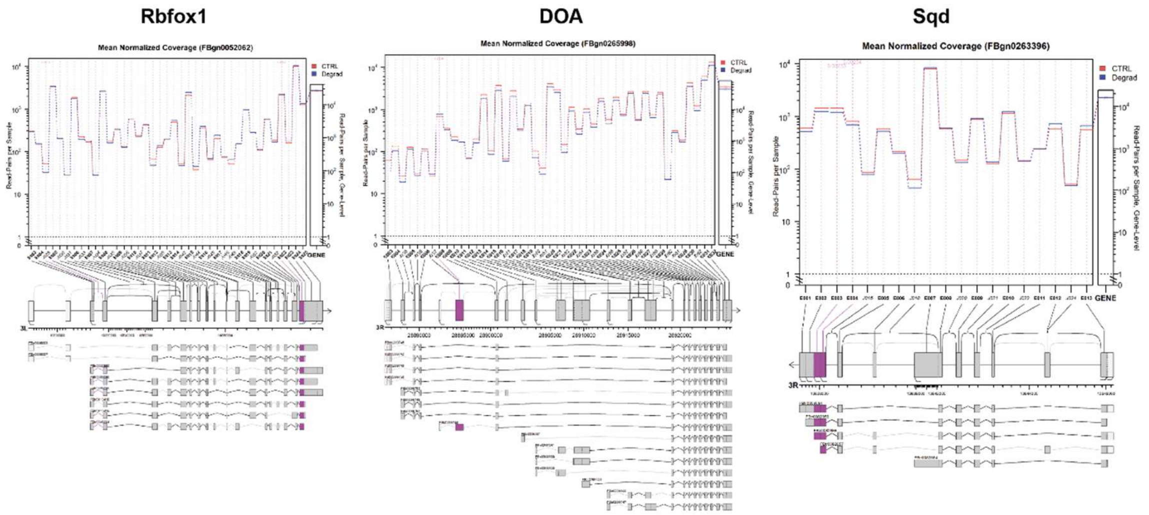


**Supplementary Figure 3: Ubx degradation impacts on alternatively spliced genes involved in mRNA processing in embryonic mesoderm.** Representative genes involved in mRNA processing (Rbfox1, DOA, Sqd) differentially spliced in embryonic mesoderm upon Ubx depletion (Domsch et al., 2019). Visualisation from Junction-Seq of the mean normalised read count for each exon or splice junction (left Y-axis, extended panel), and the gene level normalised read count (right Y-axis, narrow panel) of genes differentially spliced upon Ubx depletion in the mesoderm (degrad, blue line) compared to control (CTRL, red line). Significant differential splicing events are highlighted in purple. Isoforms including differentially spliced exon or junction usages (purple) upon Ubx degradation are displayed below each read count.

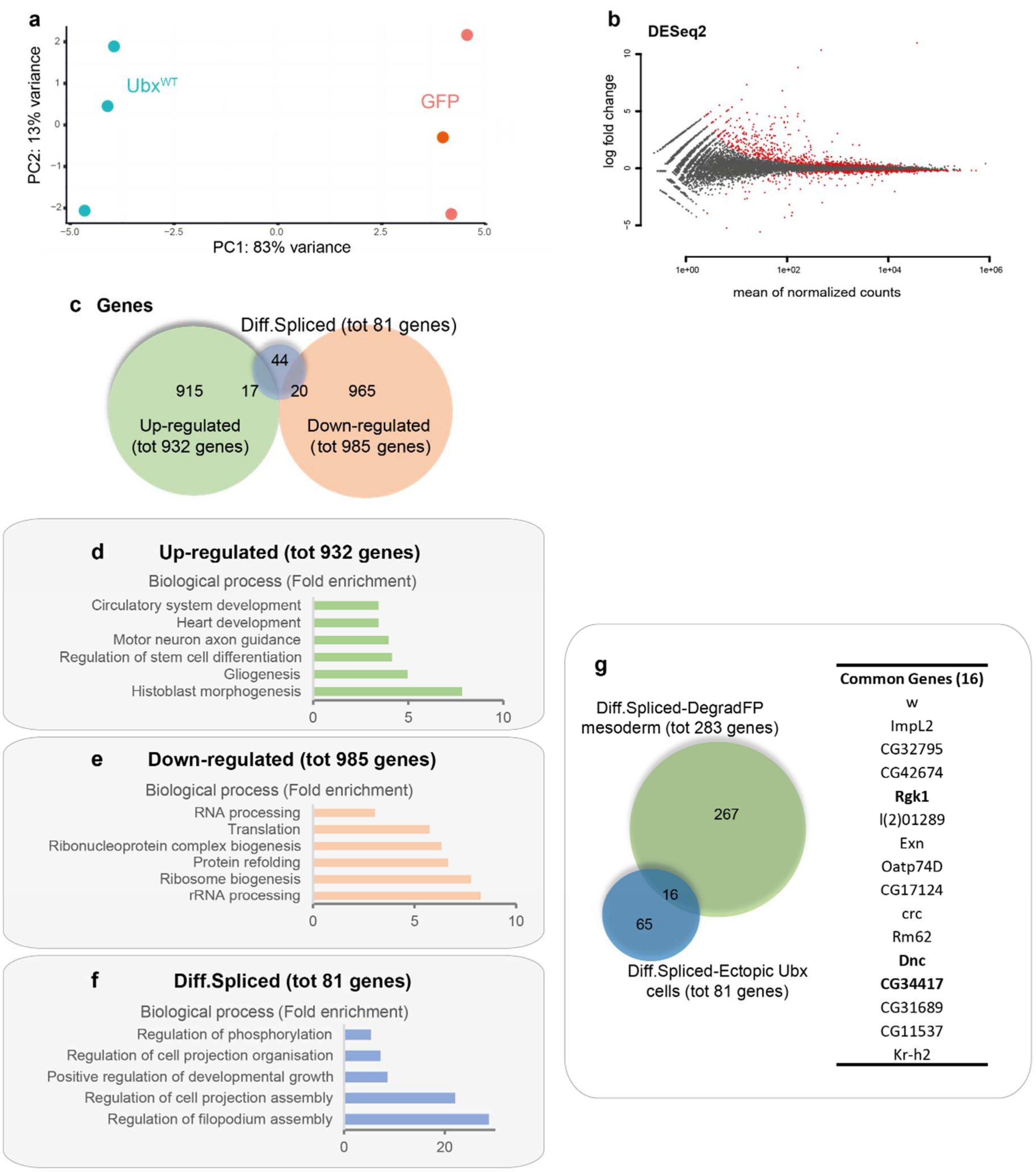


**Supplementary Figure 4: Transcriptome upon Ubx$^{WT}$ expression in *Drosophila* S2R+ cells. a.** Principle Component Analysis (PCA) applied to RNA-Seq data triplicate of *Drosophila* S2R+ expressing GFP and Ubx$^{WT}$. Two clusters corresponding to GFP and Ubx$^{WT}$ are displayed validating the replicates similarity. **b.** DEseq2 visualisation of mean normalised count and logarithmic fold change of genes up-regulated and down-regulated upon Ubx$^{WT}$ expression compared to control GFP. **c.** Venn diagram overlapping the up-regulated (932), down-regulated (985) and differentially spliced (81) genes upon Ubx$^{WT}$ expression in *Drosophila* S2R+ cells. **d-f.** Fold enrichment of Gene ontology (GO) term of biological processes enriched for the list of genes (**d**) up-regulated (932), (**e**) down-regulated (985) and (**f**) differentially spliced (81) upon Ubx$^{WT}$ expression in *Drosophila* S2R+ cells compared to

GFP (pvalue<0.05, fold enrichment). **g.** Venn diagram overlapping the genes differentially spliced upon Ubx depletion in the embryonic mesoderm (283, green) and upon $Ubx^{WT}$ expression in *Drosophila* S2R+ cells (81). The 16 genes in common are listed. The genes validated by RTqPCR in *Drosophila* S2R+ cells are emphasised in bold (Dnc, Rgk1, CG34417).

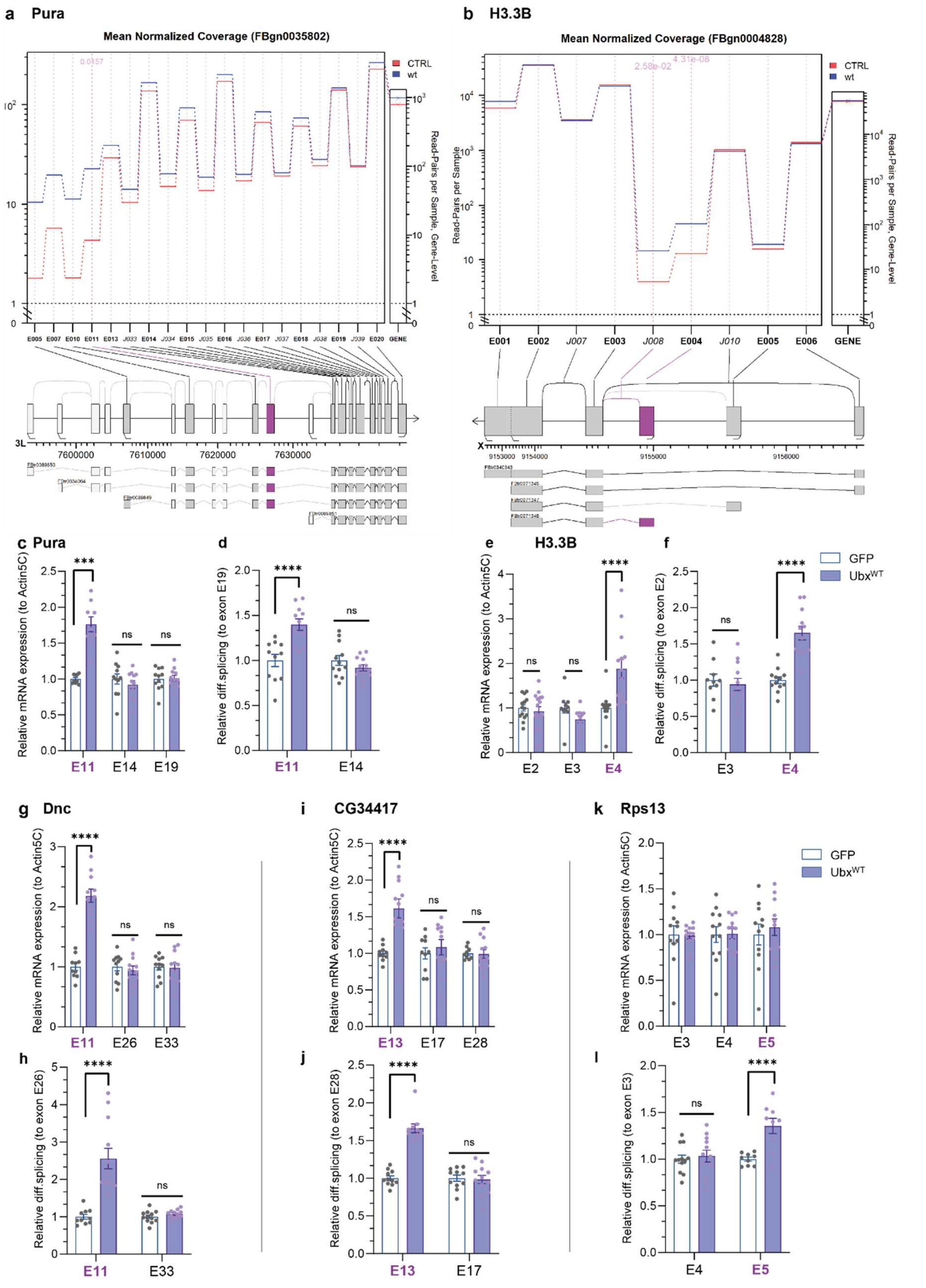

**Supplementary Figure 5: Validation of differentially spliced target genes of Ubx in S2R+ *Drosophila* cells. a-b.** Visualisation from Junction-Seq of the mean normalised read count for each exon or splice junction (left Y-axis, extended panel), and the gene level normalised read count (right Y-axis, narrow panel) of Pura (**a**) and H3.3B (**b**), differentially spliced upon Ubx$^{WT}$

expression (blue line) compared to control (red line). Significant differential splicing events are highlighted in purple. Isoforms including differentially spliced exon or junction usages (purple) upon $Ubx^{WT}$ expression are displayed bellow each read count. **c-l**. RT-qPCR experiments showing the differential expression (**c, e, g, i, k**) over actin5C and **(d, f, h, j, l**) differential retention of exon cassettes over constitutive exons for Pura (**c-d**), H3.3B (**e-f**), Dnc (**g-h**), CG34417 (**i-j**) and Rps13 (**k-l**) in cells expressing GFP control or $Ubx^{WT}$ (blue). (E+number)=exon related to Junction-Seq annotation. Differentially spliced exons are underlined (purple). n=4 independent biological triplicates. Bars represent mean±SEM. Statistical test by one-way ANOVA (p<0.001***, p<0.0001****, ns=non-significant).

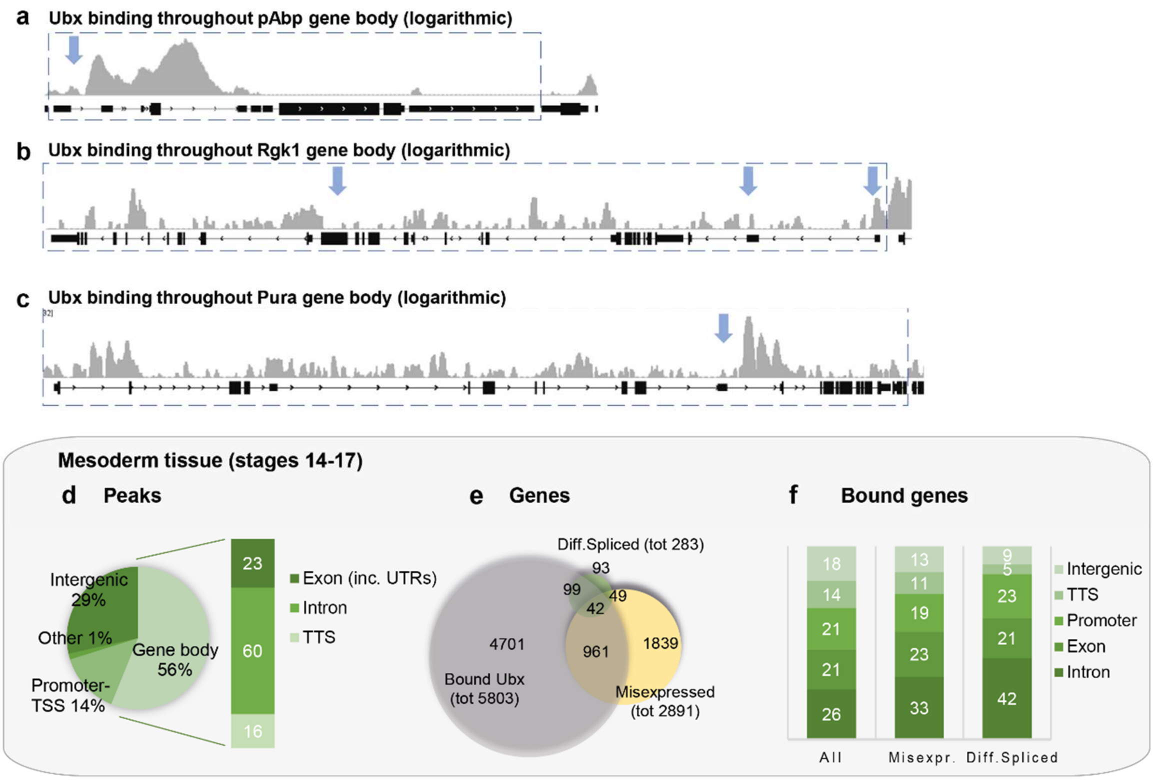


**Supplementary Figure 6: Ubx binding profiles from ChIP-Seq performed in *Drosophila* S2 cells and in the embryonic mesoderm. a-c.** Visualisation of Ubx binding events in the gene body of pAbp (**a**), Rgk1 (**b**) and Pura (**c**) by logarithmic scale in *Drosophila* S2 cells extracted from the ChIP-Seq data of Zouaz et al. (2017). **d.** Distribution of the genomic regions bound by Ubx, namely promoter and TSS (Transcription Start Site), intergenic, gene body, using the ChIP-Seq data of Ubx in embryonic mesoderm generated by Domsch et al. (2019). The distribution in the gene body is further detailed for intron, Transcription Termination Sites (TTS) and exons (including 5' and 3' UTRs). **e.** Venn diagram representing the overlap of genes bound, misexpressed and differentially spliced upon Ubx mesoderm depletion in embryos (Domsch et al., 2019). **f.** Graphical view of the distribution (%) of the total genes bound, misexpressed and bound, differentially spliced and bound by Ubx according to intergenic, promoter, exon, intron and TTS in embryonic mesoderm. Chi$^2$ tests were performed to test the distribution: "all vs misexp.": p=0.43, "all vs diff.spliced": p=0.00048, "misexp. vs diff.spliced": p=0.081.

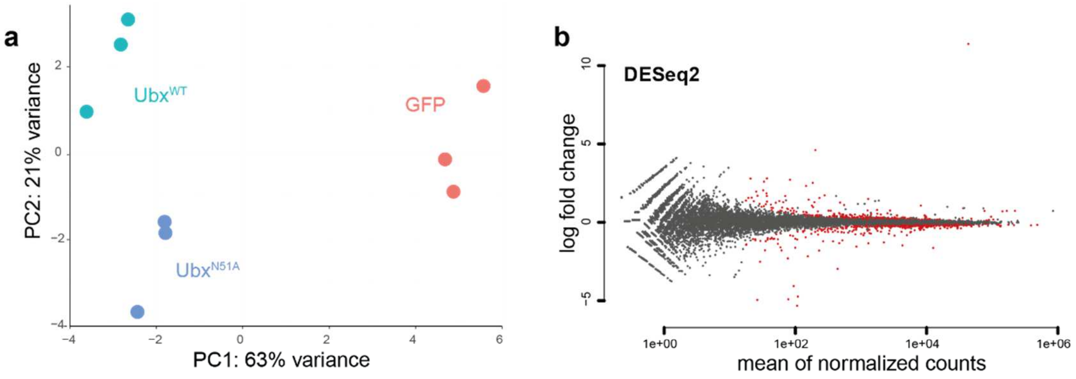


**c**

| Diff. events | Total | First exon | Higher with UbxN51A | Lower with UbxN51A |
|---|---|---|---|---|
| **Exons** | **19** | 4 | 12 | 7 |
| **Splice sites** | **5** | - | 7 | 3 |
| **Total events** | **24** | | 14 | 10 |

**d**

| | RPKM | | | | Differential expression |
|---|---|---|---|---|---|
| **Samples** | Replicate1 | Replicate2 | Replicate3 | Mean | fold change GFP |
| **Ubx$^{WT}$** | 71888 | 69408 | 102681 | 81325 | 10.98 |
| **Ubx$^{N51A}$** | 51719 | 76367 | 145254 | 91113 | 11.38 |

**Supplementary Figure 7: Transcriptome profiles in *Drosophila* S2R+ cells upon ectopic expression of Ubx$^{N51A}$. a.** Principle Component Analysis (PCA) applied to RNA-Seq data triplicate of *Drosophila* S2R+ expressing GFP, Ubx$^{WT}$ and Ubx$^{N51A}$. Three clusters corresponding to GFP, Ubx$^{WT}$ and Ubx$^{N51A}$ are displayed validating the replicates similarity. **b.** DEseq2 visualisation of mean normalised count and logarithmic fold change of genes up-regulated and down regulated upon Ubx$^{N51A}$ expression compared to control GFP. **c.** Summary of differential splicing events detected upon Ubx$^{N51A}$ expression (exons, splice sites), higher or lower compared to control. **d.** Summary of Ubx RNA expression level detected in *Drosophila* S2R+ cells upon expression of Ubx$^{WT}$ and Ubx$^{N51A}$ in each replicate. Values are indicated for RPKM, mean of the replicates and fold changes over GFP control. The values show that Ubx expression is similar for each replicate.

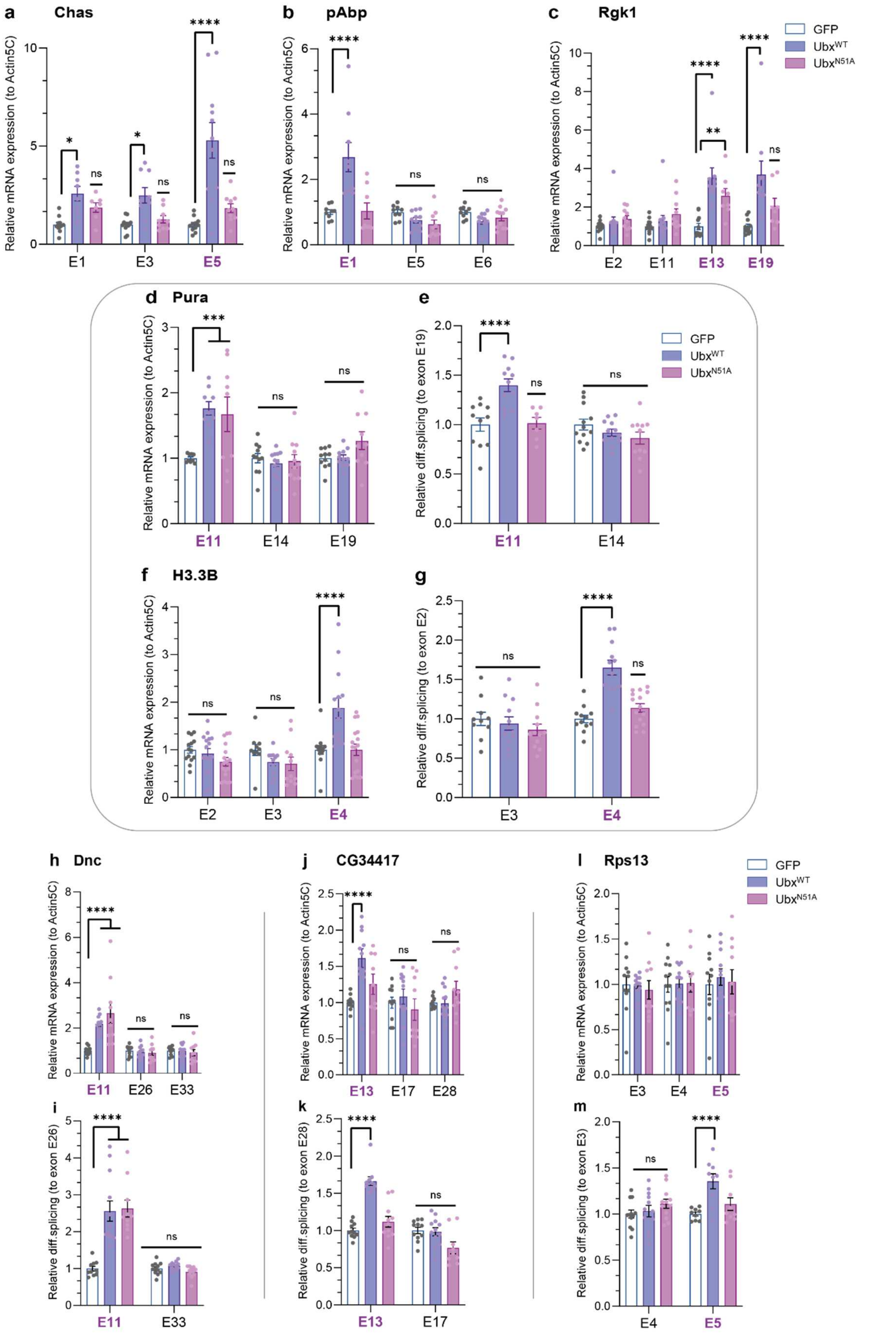


**Supplementary Figure 8: Validation of differentially spliced target genes in *Drosophila* S2R+ cells upon expression of Ubx$^{WT}$ or Ubx$^{N51A}$. a-m.** RT-qPCR experiments showing the differential expression (**a, b, c, d, f, h, i, j**) over actin5C and differential retention of exon cassettes (**e, g, k, l, m**) over constitutive exons for Chas (**a**), pAbp (**b**), Rgk1 (**c**), Pura (**d-e**), H3.3B (**f-g**), Dnc (**h, k**), CG34417 (**l, l**) and Rps13 (**j, m**) in *Drosophila* S2R+ cells expressing GFP control, $Ubx^{WT}$ (blue) or $Ubx^{N51A}$ (purple). Results showed that only Dnc is differentially spliced upon expression of $Ubx^{N51A}$ while all genes are differentially spliced exclusively upon $Ubx^{WT}$ expression. (E+number)=exon related to Junction-Seq annotation. Differentially spliced exons are underlined (purple). n=4 independent biological triplicates. Bars represent mean±SEM. Statistical test by one-way ANOVA (p<0.05*, p<0.01**, p<0.001***, p<0.0001****, ns=non-significant).

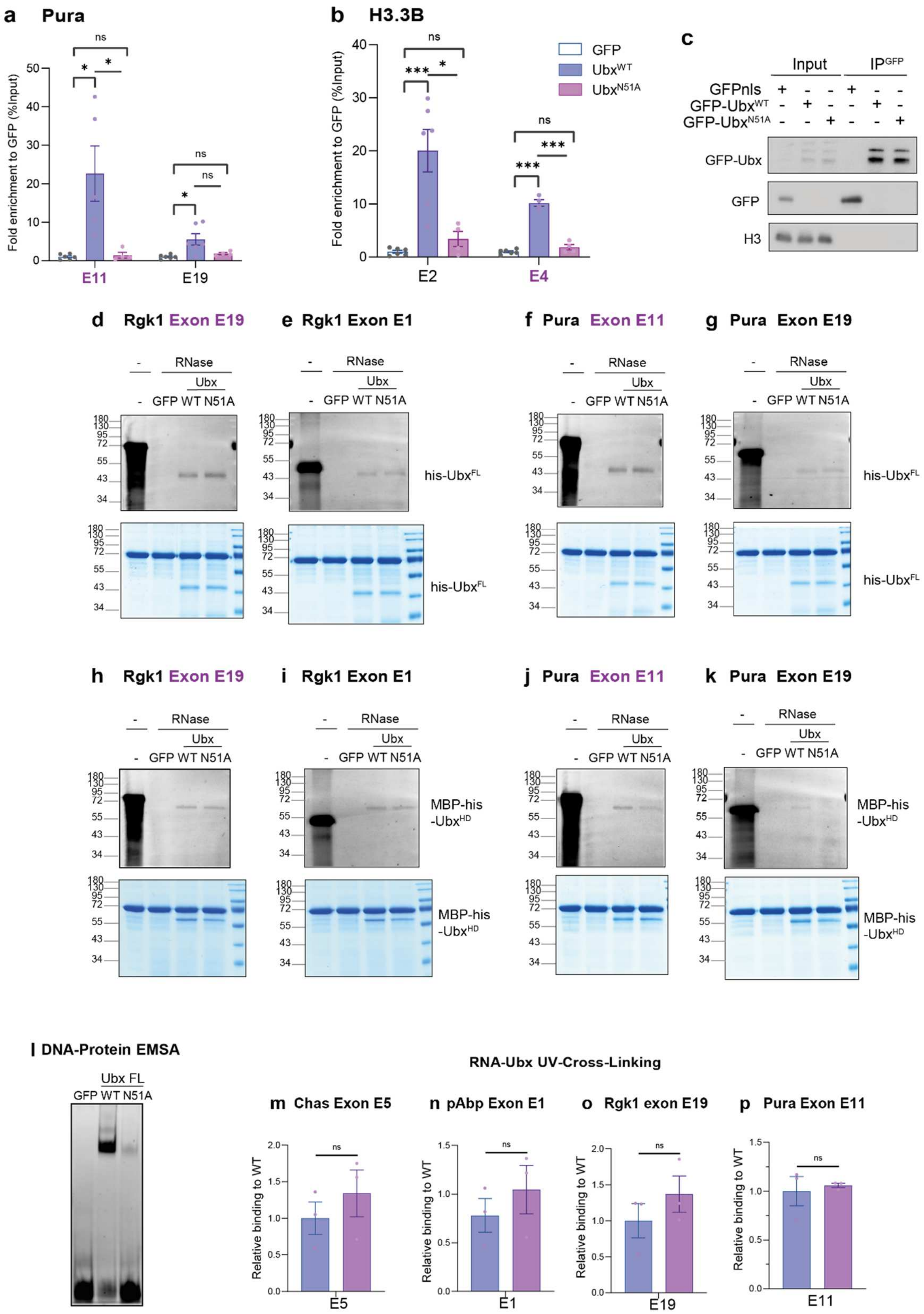


**Supplementary Figure 9: Ubx binds RNA *in vivo* and *in vitro.* a-b.** RNA-Immunoprecipitation (RIP-RTqPCR) experiments of *Drosophila* S2R+ cells expressing GFP control, GFP-Ubx$^{WT}$ (blue) and GFP-Ubx$^{N51A}$ (purple) showing an enrichment of targeted exonic regions of Pura (**a**) and H3.3B (**b**). Values are RNA relative enrichment over GFP calculated as percentage of input. (E+number)=exon related to Junction-Seq annotation, differentially spliced exons are underlined (purple). n=3 independent biological duplicates. Bars represent mean±SEM. The results exhibited a specific enrichment of differentially spliced exonic sequences in $Ubx^{WT}$ fraction compared to GFP and $Ubx^{N51A}$. **c.** Immunoblotting of RIP experiments showing the immunoprecipitation of GFP fusion proteins (GFPnls, GFP-$Ubx^{WT}$, GFP-$Ubx^{N51A}$), ectopically expressed in *Drosophila* S2R+ cells. Western blots were probed with the indicated antibodies. The input is shown as a control of expression level (lanes 1-3), Histone 3 (H3) is used as a loading control. **d-k.** Fluorescent protein-RNA interaction assay followed by UV-crosslinking and RNase digestion, performed *in vitro* with purified proteins, namely MBP-his-GFP as control, his-Ubx (WT and N51A) full length (FL) (**d-g**) and the homeodomain alone MBP-Ubx-HD (WT and N51A) (**h-k**) for Rgk1 exon E19, E1 (**d-e, h-i**) and Pura E11, E19 (**e-f, j-k**). Interactions were detected on denaturating gels by Cy3-UTP signal (upper panel), and gels were stained by Coomassie to reveal the protein content (lower panel). Each probe is indicated relative to the genes and exons. Molecular marker is indicated. Notably, MBP-his-GFP loaded samples showed a band at the same size as the BSA staining. **l.** DNA-Electrophoretic Mobility Shift Assay (EMSA) of his-fused proteins showing a binding of $Ubx^{WT}$ and a loss of DNA-binding ability for $Ubx^{N51A}$ protein. **m-p.** Graphical view showing the quantification of relative RNA-binding of $Ubx^{N51A}$ compared to $Ubx^{WT}$ protein for exon cassette Chas E5 (**m**), pAbp E1, (**n**), Rgk1 E19 (**o**), Pura E11 (**p**). Notably $Ubx^{N51A}$ conserves the exact same RNA-binding abilities as $Ubx^{WT}$. n=3 independent biological replicates. Statistical test by one-way ANOVA ($p<0.05$ *, $p<0.001$***, ns=non-significant). Differentially spliced exons are underlined (purple). See also Supplementary Table 7.

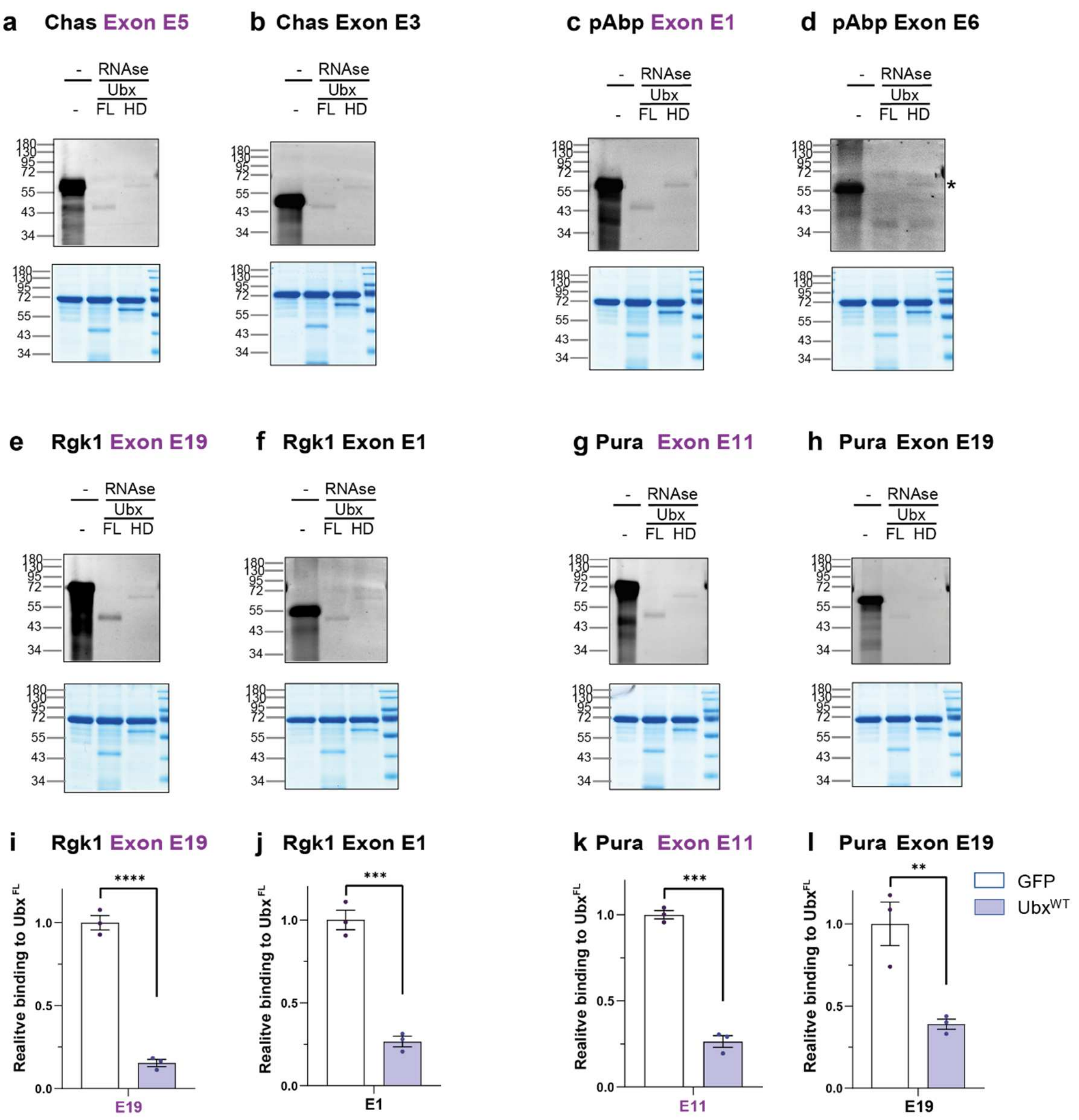


**Supplementary Figure 10: Ubx HD partly mediates RNA-binding *in vitro*. a-d.** Fluorescent protein-RNA interaction assay followed by UV-crosslinking and RNase digestion, performed *in vitro* with purified proteins his-Ubx full length (FL) and MBP-Ubx-HD (WT) for Chas exon cassette E5 (**a**), constitutive E3, (**b**), pAbp exon cassette E1 (**c**), constitutive E6 (**d**) illustrating the quantification of Fig.4. Interactions were detected on denaturating gels by Cy3-UTP signal (upper panel), and gels were stained by Coomassie to reveal the protein content (lower panel). Each probe is indicated relative to the genes and exons. Molecular marker is indicated. Notably, MBP-his-GFP exhibited a band at the same size as the BSA staining **e-h.** Fluorescent protein-RNA interaction assay followed by UV-crosslinking and RNase digestion, performed *in vitro* with purified proteins his-Ubx full length (FL) and MBP-Ubx-HD (WT) for Rgk1 exon cassette E19 (**e**), constitutive E1, (**f**), Pura exon cassette E11 (**g**), constitutive E19 (**h**) illustrating the quantifications (**i-l**). n=3 independent biological replicates. Statistical test by one-way ANOVA ($p<0.05$ *, $p<0.001$***, ns=non-significant). Differentially spliced exons are underlined (purple). See also Supplementary Table 7.

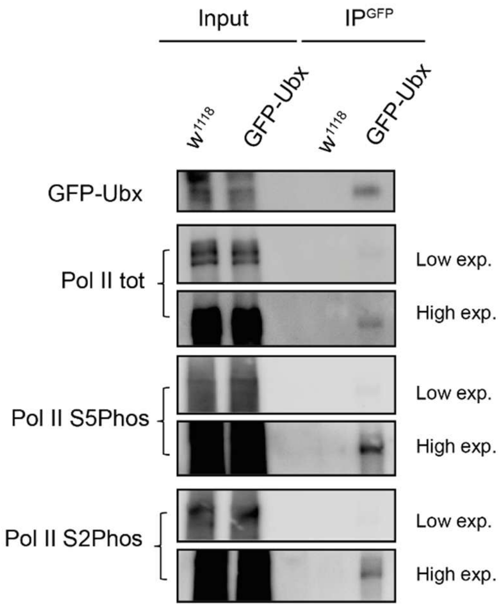


**Supplementary Figure 11: Ubx interacts with paused and active Pol II in *Drosophila* embryos.** Co-immunoprecipitation of Pol II total, paused (S5Phos) and active (S2Phos) from nuclear extract of control ($w^{1118}$) or *GFP-Ubx* embryos, which carry a CRISPR/Cas9 engineered version of the *Ubx* gene, *GFP-Ubx*, at the endogenous locus (Domsch et al., 2019). The input fraction is present as control (lane 1-2). Pol II total, S5Phos and S2Phos were co-immunoprecipitated with GFP-Ubx ($IP^{GFP}$-lane 4), but not in purified extracts from $w^{1118}$ embryos ($IP^{GFP}$-lane 3).

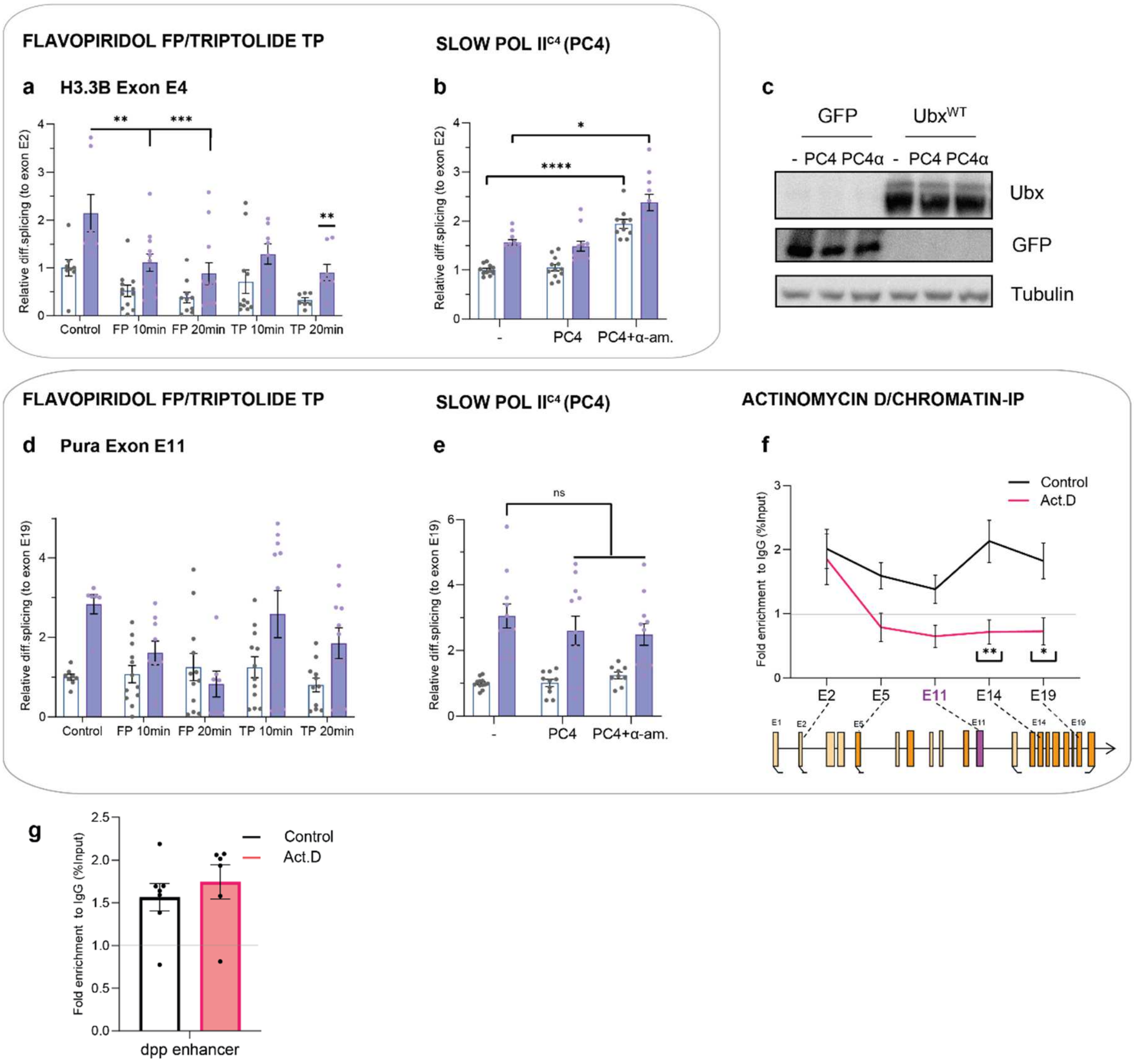


**Supplementary Figure 12: Ubx regulates splicing in an elongation-coupled manner. a-b.** RT-qPCR experiments showing the differential retention of exon cassettes over constitutive exons for H3.3B (**a**), Pura (**b**) in *Drosophila* S2R+ cells expressing GFP control, $Ubx^{WT}$ (blue) and treated for 10 or 20 minutes with Flavopiridol (FP, elongation repressed) or Triptolide (TP, elongation proceeds until termination). **c-d.** RT-qPCR experiments showing the differential retention of exon cassettes over constitutive exons for H3.3B (**c**), Pura (**d**) in cells expressing GFP control, $Ubx^{WT}$ (blue) and the slow Pol $II^{C4}$ (PC4) mutant, combined with α-amanitin treatment (α-am) showed that Ubx effect on splicing depends on the Pol II rate. **e.** Immunoblotting showing the expression level in cells expressing GFP control, $Ubx^{WT}$ and the slow Pol $II^{C4}$ (PC4) mutant, showing that PC4 expression does not affect the expression level of Ubx. Western blot was probed with the indicated antibodies. Tubulin is used as a loading control. **f-g.** Chromatin-IP experiments of Ubx coupled with actinomycin D treatment are presented as percent of enrichment relative to input and IgG control (horizontal bar set to 1). Bindings of Ubx on the proximal and distal exons to the Transcription Start Site (TSS) of Pura (**f**) are displayed relative to a schematic of the gene architecture. Binding of decapentaplegic (dpp) intergenic enhancer by Ubx is showed as control (**g**). Differentially spliced exons are highlighted in purple. n=3 independent biological in triplicates (RNA) or duplicates (ChIP). Bars represent mean±SEM. Statistical test by one-way ANOVA ($p<0.05$*, $p<0.01$**, $p<0.001$***, ns=non-significant).

**a**

| | AIC | | BIC | |
|---|---|---|---|---|
| **Sample** | **Single exp.** | **Double exp** | **Single exp.** | **Double exp** |
| GFP-Ubx$^{WT}$ | -241 | **-298** | -234 | **-285** |
| GFP-Ubx$^{WT}$+PC4 | -236 | **-284** | -228 | **-271** |

**b**

| **Sample** | **Residence time (sec)** | **Stat.** |
|---|---|---|
| GFP-Ubx$^{WT}$ | 25.2 | ns |
| GFP-Ubx$^{WT}$+PC4 | 18.6 | ns |
| GFP-Ubx$^{N51A}$ | 14.4 | ns |
| GFP-Ubx$^{N51A}$+PC4 | 20.6 | ns |

**c**

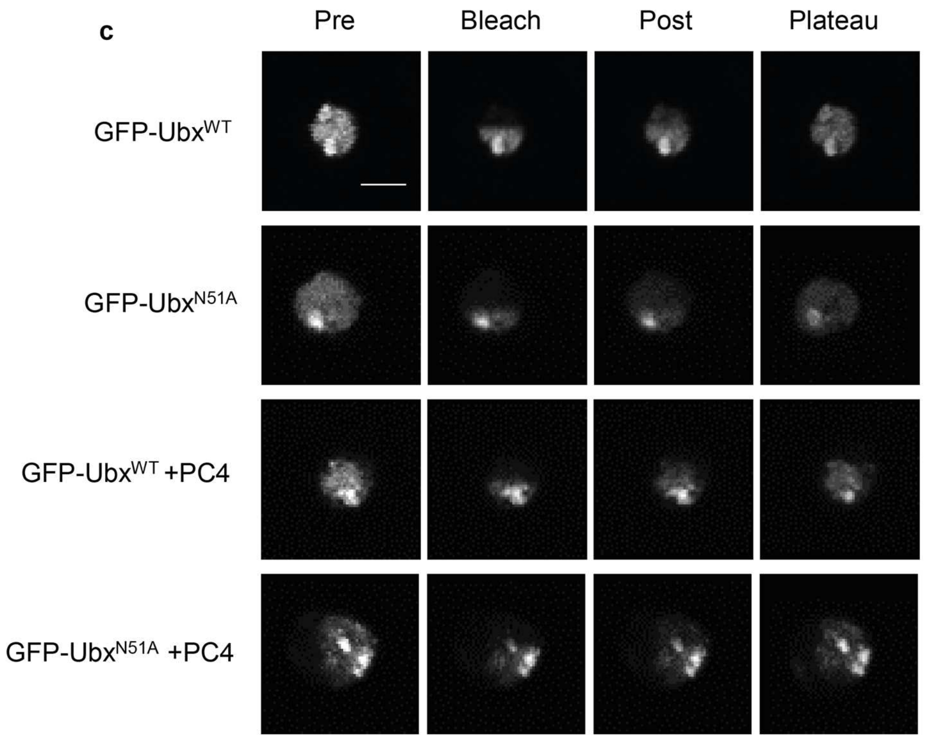


**Supplementary Figure 13: Fluorescence Recovery after Photobleaching of Ubx$^{WT}$ and Ubx$^{N51A}$ coupled with slow Pol II$^{C4}$. a.** Table of Akaike Information Criterion (AIC) and Bayesian Information Criterion (BIC) mean number related to the modelling with a single and a double exponential models, for each sample expressing GFP-Ubx$^{WT}$, GFP-Ubx$^{N51A}$ coupled with slow Pol II$^{C4}$ (PC4) and α-amanitin in *Drosophila* S2R+ cells, indicating that the double exponential model fits the experimental data (lowest number in bold). **b.** Table of mean of residence time for GFP-Ubx$^{WT}$, GFP-Ubx$^{N51A}$ coupled with slow Pol II$^{C4}$ (PC4) and α-amanitin expressing S2R+ cells. While the residence time of the Ubx$^{N51A}$ is lower than the Ubx$^{WT}$, the statistical analysis does not reflect the difference of protein dynamic, as expected due to the loss of RNA-binding abilities of the Ubx$^{N51A}$. Similarly, the slow Pol II$^{C4}$ does not significantly affect the residence time of Ubx (WT and N51A) while the half time recovery was significantly modified (see Fig.7). **c.** Imaging data of GFP-Ubx$^{WT}$, GFP-Ubx$^{N51A}$ coupled with slow Pol II$^{C4}$

(PC4) and α-amanitin in S2R+ cells. Pre-bleach, first post-bleach, 1/2recovery time and reached plateau corresponded acquisition images are shown.

**DNA binding domain alignment**

```
SP|P83949|UBX_DROME  RRRGRQTYTRYQTLELEKEFHTNHYLTRRRRIEMAHALCLTERQIKIWFQNRRMKLKKEI 60
SP|P09081|BCD_DROME  PRRTRTTFTSSQIAELEQHFLQGRYLTAPRLADLSAKLALGTAQVKIWFKNRRRRHKIQS 60
SP|P48431|SOX2_HUMAN -------------------------------------VKRPMNAFMVWSRGQRRKMAQEN 23
SP|P48436|SOX9_HUMAN -------------------------------------VKRPMNAFMVWAQAARRKLADQY 23
                                                          :      . :* :  * :    :

SP|P83949|UBX_DROME  ----------------------------------------------
SP|P09081|BCD_DROME  ----------------------------------------------
SP|P48431|SOX2_HUMAN PKMHNSEISKRLGAEWKLLSETEKRPFIDEAKRLRALHMKEHPDYK 69
SP|P48436|SOX9_HUMAN PHLHNAELSKTLGKLWRLLNESEKRPFVEEAERLRVQHKKDHPDYK 69
```

**Supplementary Figure 14: Alignment of DNA-RNA Binding Proteins and conserved domains.** Alignment of the RNA-binding domains of Ubx (HD), Bcd (HD), Sox2 (HMG) and Sox9 (HMG) showing similarity between the first helix of Sox TFs and the third helix of HD TFs. The Asparagine N51 of Ubx is highlighted in blue. The amino acid R54 of Bcd involved in *Caudal* RNA-binding is underscored. Notably, Sox and HD TFs belong to the super family of helix loop helix (HLH) TF containing RNA-binding domain (Auer et al., 2020). *: amino acids conserved; . and : are amino-acids conserved based on their similar nature

| Sequence name | Nucleotide sequence | Length | Number of U |
|---|---|---|---|
| Chas Exon 3 | CUCCAAGUACUUCCAGGGCAUCUCGAACUGGUCGGAACG CAAGGAUCCCGGUCCGCCGGUCAAUCACUUCUCGUCGCA GG | 80bp | 16 |
| Chas Exon 5 | CAAAAACCCAUAGCAGCUGGCUUUCGGCCCGGGGCAGAC CUGUUCUGAGCCAGGUUUAAUCAAUAGCCAAAGAGCUACU GCUGCUGCUGCUGCUGCUGCUGCUCCUGCUUGGCUUU | 116bp | 30 |
| pAbp Exon 1 | UCCUUGAAUGUGGUCUAUUUCGUAAAAACAAAAACACAAA AAAAAAACUUAAAAAAUUCUUCAAUCUCUGUAGUGCCGAU UUCCAUCGAGAACGAAAAGAAGUACUCAAGCAC | 113bp | 28 |
| pAbp Exon 6 | CGAGCGUCUGUAUCCGAUGAUUGAGCAUAUGCACGCCAA UUUGGCUGGUAAGAUCACCGGCAUGUUGUUGGAGAUCGA AAACUCUGAGCUCUUGCACAUGAUCGAGGA | 108bp | 28 |
| Rgk1 Exon 1 | CAGACUCGACUACUACCGGGGGUAUUAUUCCCAACUGUA CAUAUUGUGUAUAAUUUUAGUCUUAAUUUACGCUAGUCAC UUCCUCCAU | 88bp | 32 |
| Rgk1 Exon 19 | GUAAUUUCCUCGGAGCGACGACAACUUUGAAACUUGUAAA CCUUGCGGCUUGCAAUUGCAAACUGUGCAGCAAGGCGCC UAAUGUCCUCAGAUUAAAGUGCAAUCUAAUUGAUUGGCCA AGGAGCAGUGCGUGGAUCAGAGGUGCAACUAGGAGCUA | 157bp | 38 |
| Pura Exon 11 | GGAUUCGCAGCUACACAGUCUGCGCAAGAUGGGUGCUGG CCAGUUGGCCAGGCUGCAGGAUUUGGCCAGGGCCAUCAC CUCUGCUCCCGGCUCGCCGCGCAUGACGUCACAGCCGCA AUCAUCCUCAUC | 129bp | 25 |
| Pura Exon 19 | CCUCCUCGGGUCACAAUGAUGGCGGUGGCAGGCACAGCC UGCACUUUGGCAAGCUGCCUGGAGCACACAGCGCAAGUU CCACCAACUGCAGUGCCACUUUGGAGCUGAUCAACGAAAC UCAGGCCC | 126bp | 22 |

**Supplementary Table 7: Probes used for Ubx-RNA UV-crosslinking assay**

**Supplementary Table 1: differentially expressed genes in the mesoderm upon Ubx degradation**

**Supplementary Table 2: differentially spliced genes in the mesoderm upon Ubx degradation**

**Supplementary Table 3: differentially expressed genes in the *Drosophila* S2R+ cells upon Ubx$^{WT}$ ectopic expression**

**Supplementary Table 4: differentially spliced genes in the *Drosophila* S2R+ cells upon Ubx$^{WT}$ ectopic expression**

**Supplementary Table 5: differentially expressed genes in the *Drosophila* S2R+ cells upon Ubx$^{N51A}$ ectopic expression**

**Supplementary Table 6: differentially spliced genes in the *Drosophila* S2R+ cells upon Ubx$^{N51A}$ ectopic expression**

**Supplementary Table 7: Probes used for RNA-Ubx UV-crosslinking assay**

**Supplementary Table 8: Primers used in the study**
